# Transcription factor TAp73 and microRNA-449 complement each other to support multiciliogenesis

**DOI:** 10.1101/273375

**Authors:** Merit Wildung, Tilman Uli Esser, Katie Baker Grausam, Cornelia Wiedwald, Larisa Volceanov-Hahn, Dietmar Riedel, Sabine Beuermann, Li Li, Jessica Zylla, Ann-Kathrin Guenther, Magdalena Wienken, Evrim Ercetin, Zhiyuan Han, Felix Bremmer, Orr Shomroni, Stefan Andreas, Haotian Zhao, Muriel Lizé

**Affiliations:** Molecular & Experimental Pneumology Group, Clinic for Cardiology and Pneumology, University Medical Center Goettingen, Germany; Cancer Biology and Immunotherapeutics Group, Sanford Research, Sioux Falls, South Dakota, USA; Division of Basic Biomedical Sciences, University of South Dakota, Sanford School of Medicine, Vermillion, South Dakota; Electron Microscopy, Max-Planck-Institute for Biophysical Chemistry, Goettingen, Germany; Department of Genes and Behavior, MPI for Biophysical Chemistry, Goettingen, Germany; Institute of Molecular Oncology, University Medical Center Goettingen, Germany; Department of Biomedical Sciences, New York Institute of Technology College of Osteopathic Medicine, Old Westbury, New York, USA; Institute of Pathology, University Medical Center Goettingen, Goettingen, Germany; Microarray and Deep-Sequencing Core Facility, University Medical Center Goettingen, Germany

## Abstract

Motile cilia serve vital functions in development, homeostasis and regeneration. We recently demonstrated that TAp73 is an essential transcriptional regulator of respiratory multiciliogenesis. Here, we show that TAp73 is expressed in multiciliated cells (MCCs) of diverse tissues. Analysis of *TAp73* mutant animals revealed that TAp73 regulates *Foxj1, Rfx2, Rfx3*, axonemal dyneins *Dnali1* and *Dnai1*, plays a pivotal role in the generation of MCCs in male and female reproductive ducts, and contributes to fertility. However, the function of MCCs in the brain appears to be preserved despite the loss of *TAp73*, and robust activity of cilia-related networks is maintained in the absence of *TAp73*. Notably, *TAp73* loss leads to distinct changes in ciliogenic microRNAs: *miR34bc* expression is reduced, whereas the *miR449* cluster is induced in diverse multiciliated epithelia. Among different MCCs, choroid plexus (CP) epithelial cells in the brain display prominent *miR449* expression, whereas brain ventricles exhibit significant increase in *miR449* levels along with an increase in the activity of ciliogenic E2F4/MCIDAS circuit in *TAp73* mutant animals. Conversely, E2F4 induces robust transcriptional response from *miR449* genomic regions. To address whether increased *miR449* levels in the brain maintain the multiciliogenesis program in the absence of *TAp73*, we deleted both *TAp73* and *miR449* in mice. Although loss of *miR449* alone led to a mild ciliary defect in the CP, more pronounced ciliary defects and hydrocephalus were observed in the brain lacking both *TAp73* and *miR449*. In contrast, *miR449* loss in other MCCs failed to enhance ciliary defects associated with *TAp73* loss. Together, our study shows that, in addition to the airways, TAp73 is essential for generation of MCCs in male and female reproductive ducts, whereas *miR449* and TAp73 complement each other to support multiciliogenesis and CP development in the brain.

## Introduction

Cilia are hair-like appendages protruding from the cell membrane into the surrounding environment. Solitary immotile primary cilia are a common organelle in most mammalian cells, whereas motile cilia are restricted to a subset of cell types. This subset includes multiciliated cells (MCCs) lining brain ventricles, tracheal, and bronchial epithelium as well as the epithelium of male efferent ducts (EDs) and fallopian tubes (FTs) in females [1].

Multiciliogenesis requires precise regulation of the production, transport and assembly of a large number of different structural components, a process critically dependent on a hierarchical network of transcriptional and post-transcriptional regulators [2]. Geminin Coiled-Coil Domain Containing 1 (GEMC1) [3–5] and multiciliate differentiation and DNA synthesis associated cell cycle protein (MCIDAS or Multicilin) [6–8], members of the Geminin family, are early regulators of the MCC fate, downstream of the NOTCH pathway. MCC differentiation is also regulated by post-transcriptional mechanisms including microRNAs (miRNAs). *miR-34/449* constitute a conserved family that encodes six homologous miRNAs (*miR34a, 34b, 34c, 449a, 449b*, and *449c*) from three genomic loci in vertebrates. Inhibition of the NOTCH pathway e.g. by *miR449* is required for multiciliogenesis through de-repression of the transcriptional network of *GEMC1, MCIDAS*, E2F transcription factors (*E2F4, E2F5*), forkhead box J1 (*FOXJ1*), and v-myb avian myeloblastosis viral oncogene homolog (*MYB)* [9–11]. Disturbance of the molecular circuit leads to defective multiciliogenesis and ciliopathies in the airways, reproductive tracts, and the brain [1].

Transformation related protein 73 (*Trp73*) is a member of the p53 family with distinct isoforms generated from two alternative promoters: isoforms containing the N-terminal transactivation domain (TAp73), and N-terminally truncated dominant-negative isoforms (ΔNp73). Recently, we and others showed that TAp73 is essential for airway multiciliogenesis [12,13]. Gene expression analysis and chromatin immunoprecipitation (ChIP) identified TAp73 as a critical regulator of multiciliogenesis: TAp73 acts downstream of E2F4/MCIDAS and regulates the expression of *FOXJ1, RFX2*, and *RFX3* in pulmonary tissues [12,14–17].

The FT of female reproductive tract consists of MCCs that possess hundreds of motile cilia beating in a wave-like manner which, along with musculature contraction, moves the oocyte or zygote towards the uterus [18–20]. Defects in ciliary functions may lead to ectopic pregnancies or infertility [19,21]. In the male reproductive tract, MCCs in the EDs are involved in the transport of spermatozoa from testis to epididymis (Epi), their maturation and concentration [22–25].

MCCs in the brain can be found in a single layer of ependymal cells facing the ventricles and choroid plexus (CP). The CP epithelium, a specialized secretory epithelium that secretes cerebrospinal fluid, arises from monociliated progenitors in the roof plate around embryonic day (E) 12 and undergoes multi-ciliate differentiation to form multiple primary cilia [26,27]. Ependymal cells in mice are specified around day E16 and form multiple motile cilia on the apical surface after birth to facilitate cerebrospinal fluid movement [28,29]. Defects in the ependymal and CP lineages are implicated in aging, hydrocephalus, and brain tumors [30,31].

In this study, we detected robust TAp73 expression in MCCs in diverse tissues. In reproductive ducts, *TAp73* loss leads to a profound reduction of multiciliogenesis and suppression of *TAp73*-dependent transcriptional network activity. However, MCCs in the brain maintain robust multiciliogenesis activity despite *TAp73* loss. Molecular studies revealed alterations in *miR-34/449* family members in diverse MCCs from *TAp73* mutant mice: decreased levels of TAp73 target *miR34bc* concurrent with increased expression of *miR449*. In the brain, *miR449* is highly expressed in the CP and experiences significant upregulation following *TAp73* deletion. In addition, brain ventricles but no other multiciliated tissues from *TAp73* mutant animals exhibit increased expression of E2F4, which in turn is capable of eliciting robust transcriptional response from *miR449* genomic loci, suggesting that *miR449* plays a crucial role in brain multiciliogenesis in collaboration with *TAp73*. Indeed, *miR449* loss alone results in ciliary reduction in the CP, whereas loss of both *TAp73* and *miR449* leads to a dramatic reduction of multiciliogenesis in the CP and severe hydrocephalus. Therefore, the molecular network governing MCC fate is subjected to tissue-specific feedback modulation by transcriptional and post-transcriptional mechanisms.

## Materials and Methods

### Animals

*TAp73* mutant mice with a targeted deletion of exons 2 and 3 of the *Trp73* gene were a generous gift from Dr. Tak Mak (Princess Margaret Cancer Centre, *Toronto, Canada*) [32]. *miR449* mutants were previously described [33]. Both strains were maintained in C57Bl/6 background (n8) at the animal facility of the European Neuroscience Institute Goettingen, Germany in full compliance with institutional guidelines. The study was approved by the Animal Care Committee of the University Medical Centre Goettingen and the authorities of Lower-Saxony under the number 16/2069.

### Human samples

Human epididymis samples were procured with informed consent from two patients (42 and 41 years of age, respectively). All experimental procedures were approved and performed in accordance with the requirements set forth by Ethics Committee of the University Medical Centre Goettingen (application number: 18/2/16).

### Histology and immunostaining

Paraformaldehyde-fixed, paraffin-embedded tissues were treated with heat-induced epitope retrieval using Rodent Decloaker (RD913 L, Biocare Medical, *Pacheco, CA, USA*). For immunohistochemistry, endogenous peroxidase activity was quenched with 3% H2O2 for 10 min. Tissue sections were blocked with 10% fetal calf serum (FCS) in phosphate-buffered saline (PBS) with 0.1% Triton X-100, and subsequently incubated with primary antibodies (List of antibodies is provided in **Supplementary Table 1**). Biotinylated secondary antibodies were applied for 1 h at room temperature (List of antibodies is provided in **Supplementary Table 2**), after which avidin enzyme complex and substrate/chromogen were used for color development (Vector laboratories, *Burlingame, CA, USA*). Stained tissue sections were counterstained with hematoxylin. For immunofluorescence, sections were stained with fluorescently labeled secondary antibodies (List of antibodies is provided in **Supplementary Table 2**) for 1 h at room temperature. Nuclei were counterstained with 4`, 6-Diamidin-2-phenylindol (DAPI). Histology of tissue sections was assessed by using hematoxylin (Merck, *Darmstadt, Germany*) and eosin (Carl Roth, *Karlsruhe, Germany)* staining.

### Electron microscopy

Transmission electron microscopy (TEM) was performed as previously described [12]. Briefly, murine tissue samples were fixed by immersion using 2% glutaraldehyde in 0.1 M cacodylate buffer (Science Services, *München, Germany*) at pH 7.4 overnight at 4°C. Post-fixation was performed using 1% osmium tetroxide diluted in 0.1 M cacodylate buffer. After pre-embedding staining with 1% uranyl acetate, tissue samples were dehydrated and embedded in Agar 100 (Plano, *Wetzler, Germany*). Thin tissue sections (100 nm) were examined using a Philips CM 120 BioTwin transmission electron microscope (Philips Inc., *Eindhoven, The Netherlands*) and images were taken with a TemCam F416 CMOS camera (TVIPS, *Gauting, Germany*).

### Quantification of cilia markers

Cilia were quantified using the *ImageJ* software [34]. Briefly, the region of interest was selected and a threshold was set to exclude unspecific background signals. The *Analyze Particles* tool was used to measure the area of the ciliary staining. Values were normalized to the length of the epithelia measured.

### Western blot

Samples were homogenized in RIPA buffer (20 mM TrisHCl pH 7.5, 150 mM NaCl, 9.5 mM EDTA, 1% Triton X100, 0.1% SDS, 1% sodium deoxycholate) supplemented with urea (2.7 M) and protease inhibitors (Complete Mini EDTA-free, Roche, *Basel, Switzerland*). Equal amounts of protein extracts were separated by SDS-polyacrylamide gels prior to transfer onto a nitrocellulose membrane and incubated with primary antibodies (List of antibodies is provided in **Supplementary Table 1**). The membrane was washed and incubated for 1 h with horse radish peroxidase (HRP)-conjugated secondary antibodies (List of antibodies is provided in **Supplementary Table 2**) followed by chemiluminescence detection. β-ACTIN or heat shock cognate 71 kDa protein (HSC70) were used as protein loading controls.

### RNA extraction, quantitative PCR, small RNA sequencing, and RNAscope

Tissue samples were snap-frozen in liquid nitrogen and total RNA was isolated by Extrazol (7BioScience, *Hartheim, Germany*)/Chloroform extraction followed by 80% ethanol precipitation at −20°C. For cDNA synthesis, 1 μg of total RNA was incubated with the M-MuLV reverse transcriptase and a mix of random nonameric and polyA tail primers at 42°C for 1 h in a total volume of 50 µl. All reactions were set up in triplicate with self-made SYBR Green quantitative PCR (qPCR) Mix (Tris-HCl [75 mM], (NH_4_)_2_SO_4_ [20 mM], Tween-20 [0.01% v/v], MgCl_2_ [3 mM], Triton X-100 [0.25% v/v], SYBR Green I (1:40 000), dNTPs [0.2 mM] and Taq-polymerase [20 U/ml]) using 250 nM of each gene-specific primer (List of primers is provided in **Supplementary Table 3**). Standard curve method was used to assess relative transcript content. Transcript of interests were normalized to the reference transcript of ribosomal phosphoprotein P0 (Rplp0, or *36b4*) and normalized to the mean value of control samples. The results for each sample were obtained by averaging transcript levels of technical triplicates. No RT controls and dilution curves as well as melting curves and gel electrophoresis assessment of amplicons were performed for all primer combinations. For *miR449a, miR34b*, and *miR34c* quantification, TaqMan MicroRNA Assay (Applied Biosystems, Thermo Fisher Scientific, *Waltham, MA, USA*) was performed according to the manufacturer’s instructions with U6 snRNA as internal control.

Copy number in RNA samples was determined by qPCR using a murine TAp73 plasmid (MC219984, Origene, *Rockville, USA*) with a known copy number as standard curve. Copy number of the TAp73 plasmid was determined using the following formula: number of copies = (plasmid amount [ng] * 6.022×10^23^ [molecules/mole]) / (plasmid length [bp] * 1×10^9^ [ng/g] * 650 [g/mol])

The libraries for small RNA samples were prepared using TruSeq Small RNA Library Prep Kit - Set A (24 rxns) (Set A: indexes 1-12; Cat N°: RS-200-001, Illumina, *San Diego, CA, USA*) using 1 µg of total RNA according to manufacturer’s recommendations. Samples were sequenced on the Illumina HiSeq 4000 using a 50 bp single-end approach. Mapping, prediction of novel miRNAs, quality control, and differential expression (DE) analysis were carried out using Oasis2.0 (*Oasis: online analysis of small RNA deep sequencing data*) [35]. In brief, FASTQ files were trimmed with cutadapt 1.7.1 [36] removing Truseq adapter sequences (TGGAATTCTCGGGTGCCAAGG) followed by removing sequences smaller than 15 or larger than 32 nucleotides. Trimmed FASTQ sequences were aligned to mouse small RNAs using STAR version 2.4.1d [37] with a mismatch of 5% of the sequence length and by utilizing the following databases: Mirbase version 21 for miRNAs; piRNAbank V.2 for piwiRNA; and Ensembl v84 for small nuclear RNA, small nucleolar RNA, and ribosomal RNA. Counts per small RNA were calculated using featureCounts v1.4.6 [38]. Novel miRNAs were searched for using miRDeep2 version 2.0.0.5 [39]. Differential expression of small RNA was determined by DESeq2 [40], where small RNAs were considered differentially expressed with an adjusted p-value <0.05 and absolute log2 fold-change >1. The results of the DE analysis can be found in **Supplementary Table S6**, and the small RNA-seq data sets can be found in Gene Expression Omnibus (GEO) with accession number **GSE108385**.

*TAp73* (probe no. 475741), *Mcidas* (probe no. 510401-C2), *Hes1* (probe no. 417701), and *Hes5* (probe no. 400991-C2) were visualized using RNAscope 2.5 HD Duplex Reagent Kit (#322430, Advanced Cell Diagnostics, *Hayward, CA, USA*) according to manufacturer’s instructions.

### Chromatin immuno-precipitation (ChIP)

Chromatin was harvested from Saos2 cells transiently overexpressing TAp73α, TAp73β, and the control vector pcDNA3.1. Saos2 cells were routinely tested negative for Mycoplasma. ChIP and qPCR was performed as previously described using gene specific primers (sequence information is provided in **Supplementary Table 4**) [12]. Enrichment levels were determined as the number of PCR products for each gene relative to total input.

### Luciferase assay

Luciferase assay was performed as previously described [12]. Briefly, Saos2 cells were transfected with pcDNA3.1 empty vector, or pcDNA3.1 vector carrying *E2F4* or *MCIDAS*, or both *E2F4* and *MCIDAS* vectors. Moreover, a firefly luciferase reporter construct containing the putative three wild type E2F-binding sequences of *miR449* genomic region (wild type, or “WT”), or the same sequences lacking the strongest predicted E2F-binding motif (mutant, or “Mut”) were transfected (sequence information is provided **Supplementary Table 5**). In addition, a Renilla TK luciferase vector was co-transfected. At 24 h after transfection, cells were harvested and the luciferase activities were measured using the dual luciferase assay. Firefly luciferase activities were determined relative to those of Renilla TK luciferase vector and normalized to the mean value of samples from the control vector. Luciferase assays were performed as technical triplicates on every biological replicate.

### Video microscopy

Murine fallopian tube and testis connected to the epididymis were dissected and transferred to Dulbecco’s Modified Eagle’s Medium (DMEM, Gibco, Thermo Fisher Scientific, *Waltham, MA, USA*). To image spermatozoa, the epididymis was separated from testis and vas deferens and an incision was made at distal end to release the spermatozoa. Spermatozoa as well as the peristaltic contraction of the fallopian tube were imaged with an inverse microscope.

### Imaging of cilia-generated bead-flow and cilia beating in the brain ventricular system

Mouse brains were dissected and transferred to DMEM 21063 (Gibco, Thermo Fisher Scientific). Coronal slices containing the lateral ventricle, ventral third ventricle, aqueduct, and fourth ventricle were prepared by using a coronal adult brain matrix (ASI Instruments, *Warren, MI, USA*). The ventral third ventricle was processed further as previously described [41]. Tissue explant was placed in DMEM containing fluorescent latex beads (Fluoresbrite Multifluorescent 1.0 micron Microspheres, Polysciences, *Warrington, PA, USA*). Movement of fluorescent beads along the ventricular wall and within ventricular lumen was observed by fluorescence microscopy using a DMR (Leica, *Wetzlar, Germany*) upright microscope with an epifluorescence lamp. Ciliary beating was observed by differential interference contrast microscopy using the same set-up. Bead movement was recorded using a high-speed camera (Cascade II-512, Photometrics, *Tucson, AZ, USA*) operated by MultiRecorder Software (developed by Johannes Schröder_Schetlig) and analyzed using ImageJ software [34].

### Statistical Analysis

One-tailed, unpaired Student’s *t*-test assuming normal distribution and equal variances was used to calculate statistical significance for pairwise comparisons. Luciferase assay statistics were assessed using one-way ANOVA assuming normal distribution followed by Dunnett’s multiple comparison tests. The following indications of significance were used: \**P*<0.05, \*\**P* <0.01, \*\*\**P* <0.001. N values represent biological replicates. Error bars indicate standard error of the mean (SEM).

## Results

### TAp73 is expressed in diverse multiciliated epithelia

We and others previously showed that TAp73 expressed in respiratory epithelia controls multiciliogenesis [12,13]. However, little is known about the expression and function of TAp73 in other MCCs. To address this, we performed immunostaining and *in situ* hybridization and demonstrated that in addition to the testis [42,43], TAp73 is expressed in EDs, FTs, and ependymal and CP epithelial cells in the brain (**Fig. 1a-f; Supplementary Fig. 1**). qPCR and western blot analyses showed that, among different multiciliated epithelia, FTs and EDs exhibit higher levels of TAp73 expression than testis or brain (**Fig. 1g-i**). Taken together, these results demonstrate robust TAp73 expression in different MCC types.

**Fig. 1.**
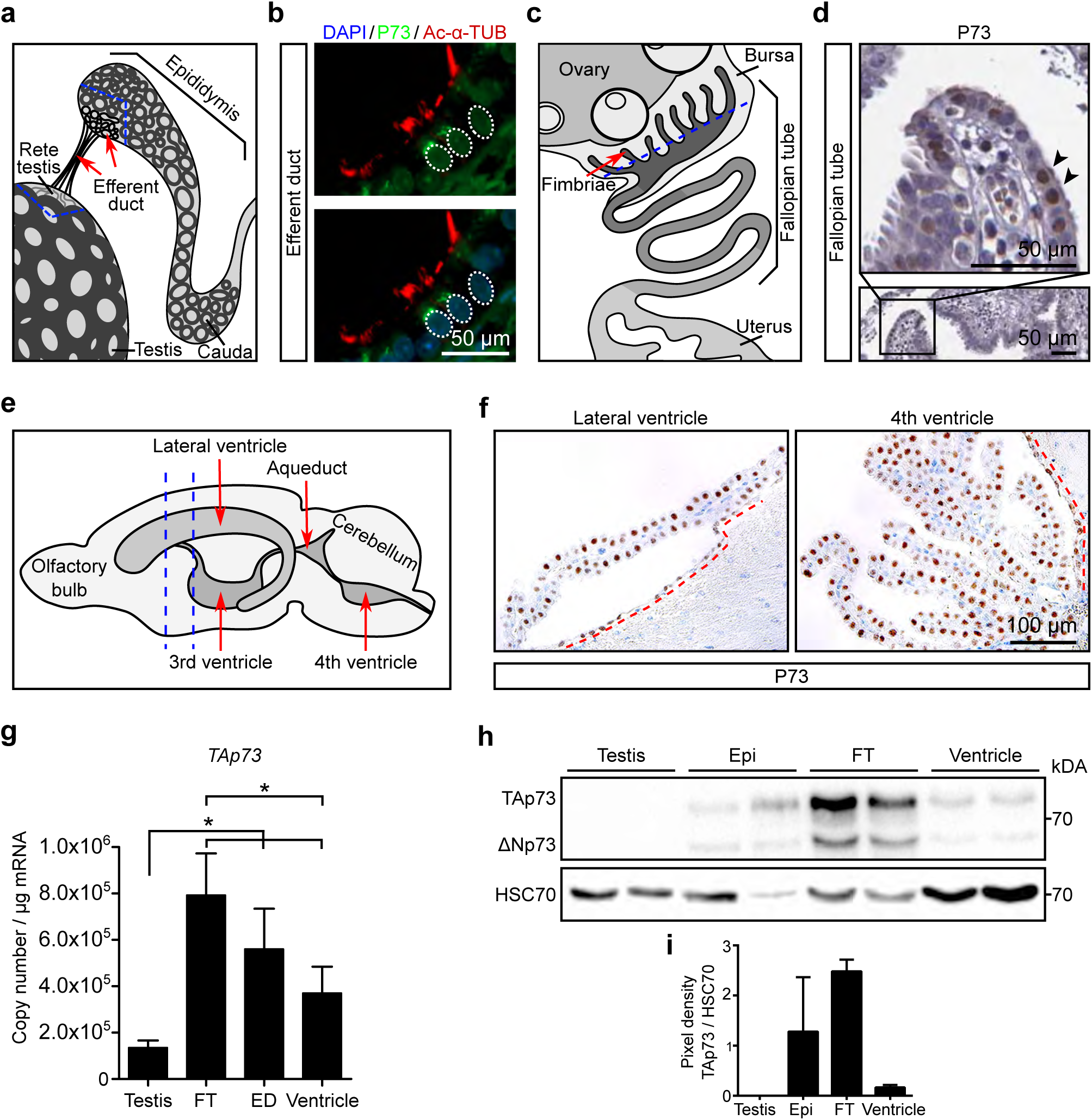
TAp73 is expressed in diverse multiciliated epithelia. (**a**) Schematic illustration of efferent ducts (EDs, red arrows) that connect testis and epididymis (Epi). Blue dotted lines indicate regions used for histological, protein, and RNA analyses. (**b**) Representative images of the expression of P73 (green) and the axonemal marker acetylated-alpha tubulin (Ac-α-TUB, red) in the human ED. White bracket circles delineate P73 nuclear staining. DAPI staining (blue) marks nuclei. (**c**) Schematic illustration of the fallopian tube (FT) that connects ovary and uterus. Blue dotted line illustrates the region used for immunofluorescence analysis including fimbriae (red arrow). (**d**) Expression of P73 in human FT. Upper panel depicts a magnification of the boxed region in the lower panel. Arrowheads mark P73^+^ cells. Images were retrieved from Human Protein Atlas (https://www.proteinatlas.org/ENSG00000078900-TP73/tissue/fallopian+tube). (**e**) Schematic illustration of murine brain ventricles (red arrows). Blue dotted lines indicate the position of coronal brain slices used in protein and RNA analyses. (**f**) Expression of TAp73 in lateral and 4^th^ ventricles of wild type (WT) adult mice. Red dotted lines demarcate ependymal cells lining brain ventricles. Notice that both ependymal and choroid plexus (CP) epithelial cells express TAp73. (**g**) Quantitative PCR analysis of *TAp73* expression in the testis, EDs, FTs, and brain ventricles from WT adult mice. Expression levels are shown in copy number. Data from a single experiment are shown (testis, *n*=3; FT and ventricle, *n*=4; ED, *n*=6). (**h**) Western blot analysis of the expression of TAp73 and ΔNp73 in testis, Epi, FT, and brain ventricle from WT adult mice. Heat shock cognate 71 kDa protein (HSC70) serves as a loading control. Data are representative of two independent experiments. (**i**) Quantitation of the signal intensity of TAp73 bands relative to that of HSC70 (**h**) is shown. All data are presented as mean ± SEM with \**P*<0.05.

### *TAp73* is crucial for the molecular circuit of multiciliogenesis in efferent ducts

Loss of *TAp73* leads to male infertility that has been attributed to defective germ cell maintenance during spermatogenesis [42,43]. Interestingly, we detected spermatozoa in testis from *TAp73* KO mice, although at a markedly reduced levels (**Supplementary Fig. 2a, b**). Despite normal morphology and mobility of these cells, no mature spermatozoa were detected in the epididymis (Epi) of these mice (**Fig. 2a; Supplementary Video 1a-d**), suggesting that additional defects may contribute to infertility. The multiciliated epithelium of the EDs contributes to gamete transport by facilitating testicular fluid circulation, fluid reabsorption, and spermatozoa concentration [22,24,25], all essential aspects of male fertility [9,44,45]. Indeed, though no gross morphological difference was observed in EDs between control and *TAp73* KO animals (**Fig. 2a**), immunofluorescent staining of the cilia components acetylated alpha-tubulin (Ac-α-TUB) and dynein axonemal intermediate chain 1 (DNAI1) showed a dramatic reduction in the number and length of cilia in the EDs from *TAp73* KO mice (**Fig. 2b, c)**. In contrast to the abundant long cilia of WT cells, mutant MCCs generated far fewer cilia as observed by transmission electron microscopy (TEM) (**Fig. 2d; Supplementary Fig. 2c**), resembling the loss of airway cilia in these animals [12]. Consistent with its role as transcriptional regulator, ChIP followed by qPCR revealed significant enrichment of TAp73 in genomic loci of *FOXJ1* [12] and dynein axonemal light intermediate chain 1 (*DNALI1)* and *DNAI1*, both encoding axonemal dyneins (**Fig. 2e; Supplementary Fig. 3**). Accordingly, expression of *Dnali1, Foxj1, Rfx2*, and *Rfx3* was reduced or almost completely lost in male reproductive ducts from *TAp73* KO animals (**Fig. 2f, g; Supplementary Fig. 2d**). Together, our data indicate that TAp73 directs *Dnali1* and *Dnai1* in addition to known critical nodes including *Foxj1, Rfx2*, and *Rfx3* to mediate multiciliogenesis in EDs (**Fig. 7a, b**). Thus, these additional defects in the multiciliated epithelium of the EDs may contribute to male infertility in *TAp73* KO mice.

**Fig. 2.**
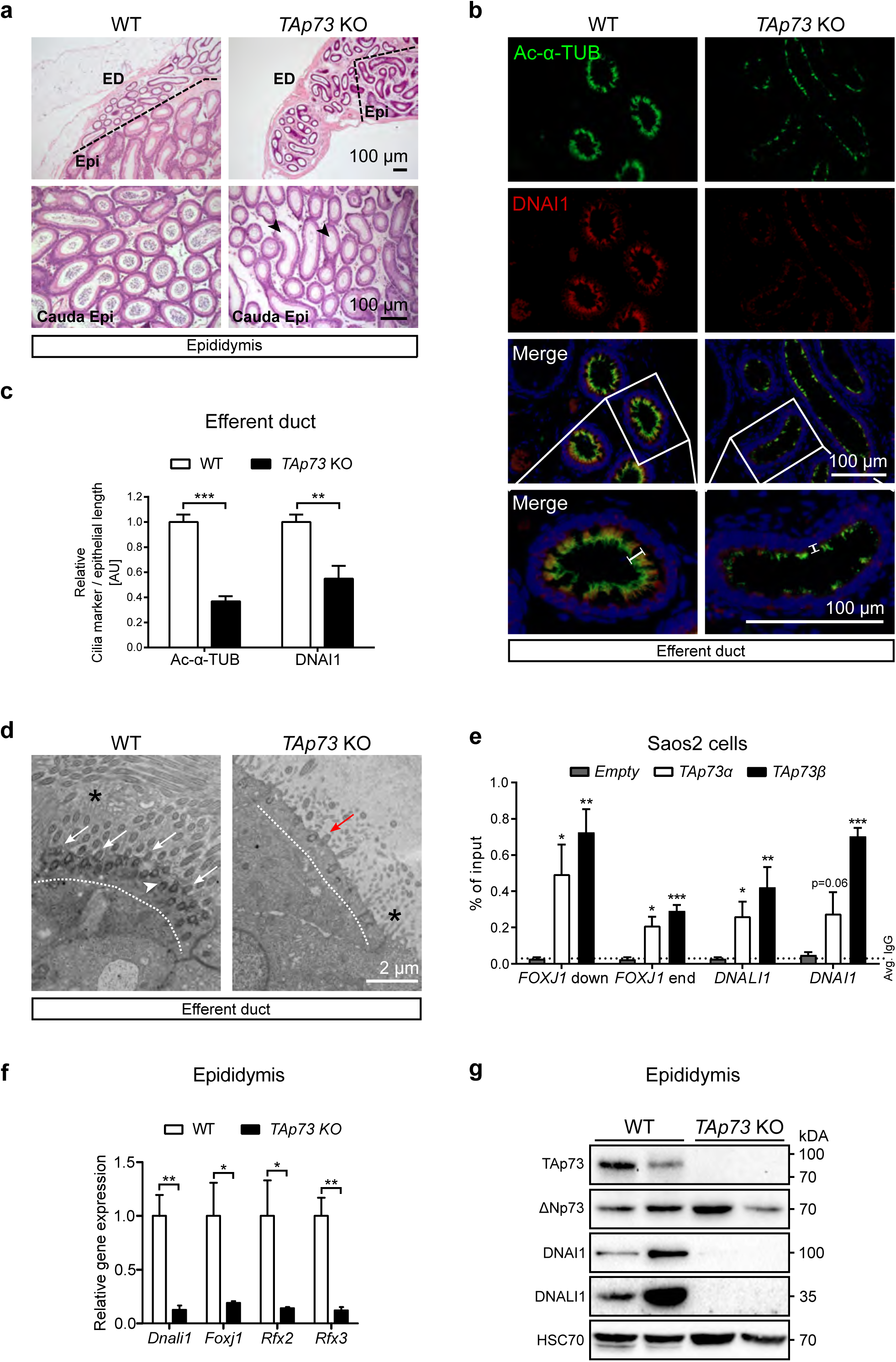
TAp73 controls multiciliogenesis in the male reproductive tract. (**a**) Representative images of hematoxylin and eosin (H&E) staining of Epi sections from WT and *TAp73* knockout (KO) animals. Bracket lines demarcate the border of EDs and Epi. Notice the lack of mature spermatozoa in cauda Epi from *TAp73* KO mice (arrowheads). (**b**) Representative images of the expression of Ac-α-TUB (green) and axonemal dynein DNAI1 (red) in EDs from WT and *TAp73* KO mice. DAPI staining (blue) labels nuclei. Boxed regions are magnified in the bottom panel. Note that *TAp73* KO mice have less cilia that also exhibit reduced length (white bars). (**c**) Quantitation of Ac-α-TUB and DNAI1 signals normalized to epithelial length. Data from a single experiment are shown (WT, *n*=6 images from 3 animals; *TAp73* KO, *n*=11 images from 4 animals). (**d**) Representative photomicrographs of transmission electron microscopy (TEM) in EDs from WT and *TAp73* KO mice. Dotted lines mark apical region of the cells. Notice the abundant cilia (white arrows) and clustered basal bodies (white arrowhead) docked to the apical surface of WT cells, whereas mutant cells exhibit fewer cilia (red arrow). Interspersed microvilli are marked with asterisks. (**e**) Chromatin immunoprecipitation was performed for Saos2 cells transfected with TAp73α, TAp73β, and empty vector. Binding of TAp73α and TAp73β to genomic regions of *FOXJ1*, axonemal dyneins *DNALI1* and *DNAI1* was evaluated by quantitative PCR and compared to vector control (*n*=3 for each antibody/gene pair, except for DNALI1 [*n*=4], genomic regions examined are illustrated in **Supplementary Fig. 3**; [76]). (**f**) Semi-quantitative PCR analysis of *Dnali1, Foxj1, Rfx2, and Rfx3* expression in EDs from WT and *TAp73* KO mice. Data from a single experiment are shown (WT, *n*=4 for *Dnali1, Foxj1*, and *Rfx3, n*=3 for Rfx2; *TAp73* KO, *n*=3). (**g**) Immunoblot analysis of the expression of TAp73, ΔNp73, DNAI1, and DNALI1 in Epi from WT and *TAp73* KO animals. HSC70 serves as a loading control. Representative result of three independent experiments is displayed. All data are presented as mean ± SEM and relative to the WT group with \**P*<0.05, ** *P* <0.01, *** *P* <0.001.

### TAp73-driven transcriptional network regulates multiciliogenesis in fallopian tubes

Though infertility in *TAp73* KO females is thought to arise from defects of oocyte development and release from the ovary [32,46], it remains unclear whether *TAp73* loss affects the multiciliated epithelium of the FT, thereby possibly influencing ova transport. Despite normal tubal morphology, analysis of Ac-α-TUB and DNAI1 expression showed reduced cilia coverage of the oviduct epithelium (**Fig. 3a-c)**. Consistently, TEM demonstrated reduced cilia and mislocated basal bodies in FTs from *TAp73* KO mice (**Fig. 3d; Supplementary Fig. 4a**). Transcript levels of *Dnali1, Foxj1*, and *Rfx2*, but not *Rfx3* were reduced in *TAp73* KO FTs (**Fig. 3e**), which was accompanied by declined protein expression of FOXJ1, DNAI1, DNALI1 (all expressed in the human FTs, **Supplementary Fig. 4b**), and gamma-tubulin (γ-TUB, basal body marker) (**Fig. 3f**), though to a lesser degree when compared to the decrease in multiciliogenesis activity observed in *TAp73*-deficient EDs. Further, smooth muscle contraction pattern in FTs is similar between control and *TAp73* KO animals (**Supplementary Video 2a, b**). Taken together, our data indicate that *TAp73* loss leads to reduced multiciliogenesis in the oviducts (**Fig. 7a, c**).

**Fig. 3.**
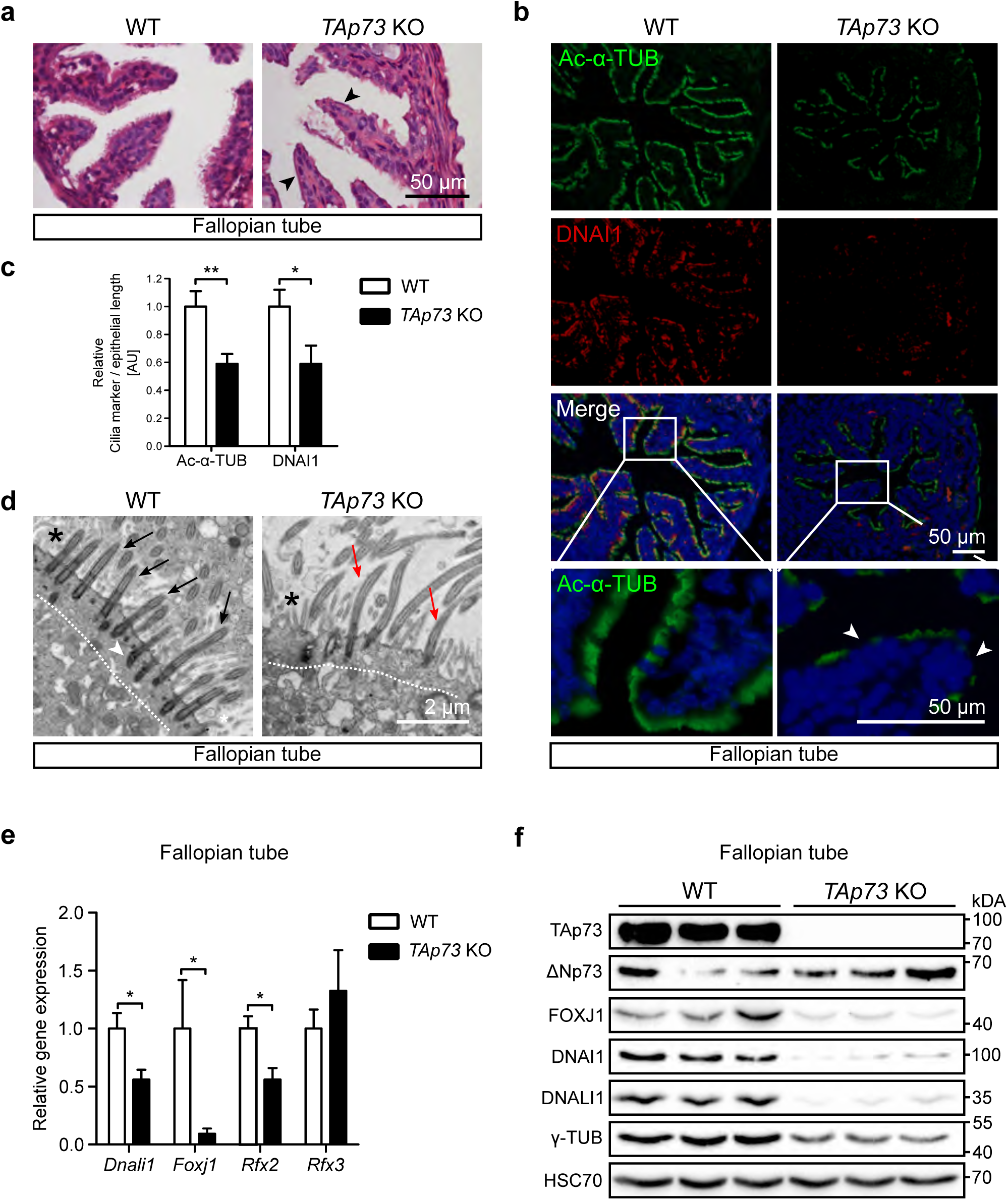
TAp73 controls multiciliogenesis in the oviducts. (**a**) Representative H&E staining of FTs from WT and *TAp73* KO animals. (**b**) The expression of Ac-α-TUB (green) and DNAI1 (red) in FTs from WT and *TAp73* KO mice. DAPI staining (blue) labels nuclei. Boxed regions are magnified in the bottom panel. In contrast to multiciliated epithelia in WT mice, *TAp73* KO mice exhibit FT segments devoid of cilia (arrowheads). (**c**) Quantitation of Ac-α-TUB and DNAI1 signals normalized to epithelial length. Data from a single experiment are shown (WT, *n*=6 images from 4 mice; *TAp73* KO, *n*=6 images from 3 mice). (**d**) Representative TEM photomicrographs of FTs from WT and *TAp73* KO animals. Dotted lines mark apical region of the cells. Notice the presence of abundant cilia (black arrows) and basal bodies (white arrowhead) docked at the apical surface of WT cells, whereas mutant cells display fewer cilia (red arrows). Interspersed microvilli are marked with asterisks. (**e**) Semi-quantitative PCR analysis of *Dnali1, Foxj1, Rfx2*, and *Rfx3* expression in oviducts from WT and *TAp73* KO mice. Data from a single experiment are shown (*n*=3). (**f**) Immunoblot analysis of the expression of TAp73, ΔNp73, FOXJ1, DNAI1, DNALI1, and gamma tubulin (γ-TUB) in oviducts from WT and *TAp73* KO animals. HSC70 serves as a loading control. Data are representative of three independent experiments. All data are presented as mean ± SEM and relative to the WT group with \**P*<0.05, ** *P* <0.01.

### Ciliary function in the brain is intact in the absence of *TAp73*

Given TAp73 expression in ependymal and CP epithelial cells, we further evaluated TAp73 expression during embryonic brain development. Immunofluorescent studies showed that proliferative progenitors (KI-67^+^) are present in hindbrain roof plate at day E14.5, whereas post-mitotic cells expressing aquaporin 1 (AQP1) [31,47] are detected in CP epithelium (KI-67^-^/AQP1^+^) (**Fig. 4a**). Notably, a portion of the roof plate exists between the progenitors and CP epithelium that remains undifferentiated after cell cycle exit (KI-67^-^/AQP1^-^) (**Fig. 4a**). In contrast to progenitors with a solitary primary cilium, the “transition” zone is comprised of MCCs that exhibit TAp73 expression (**Fig. 4b**).

**Fig. 4.**
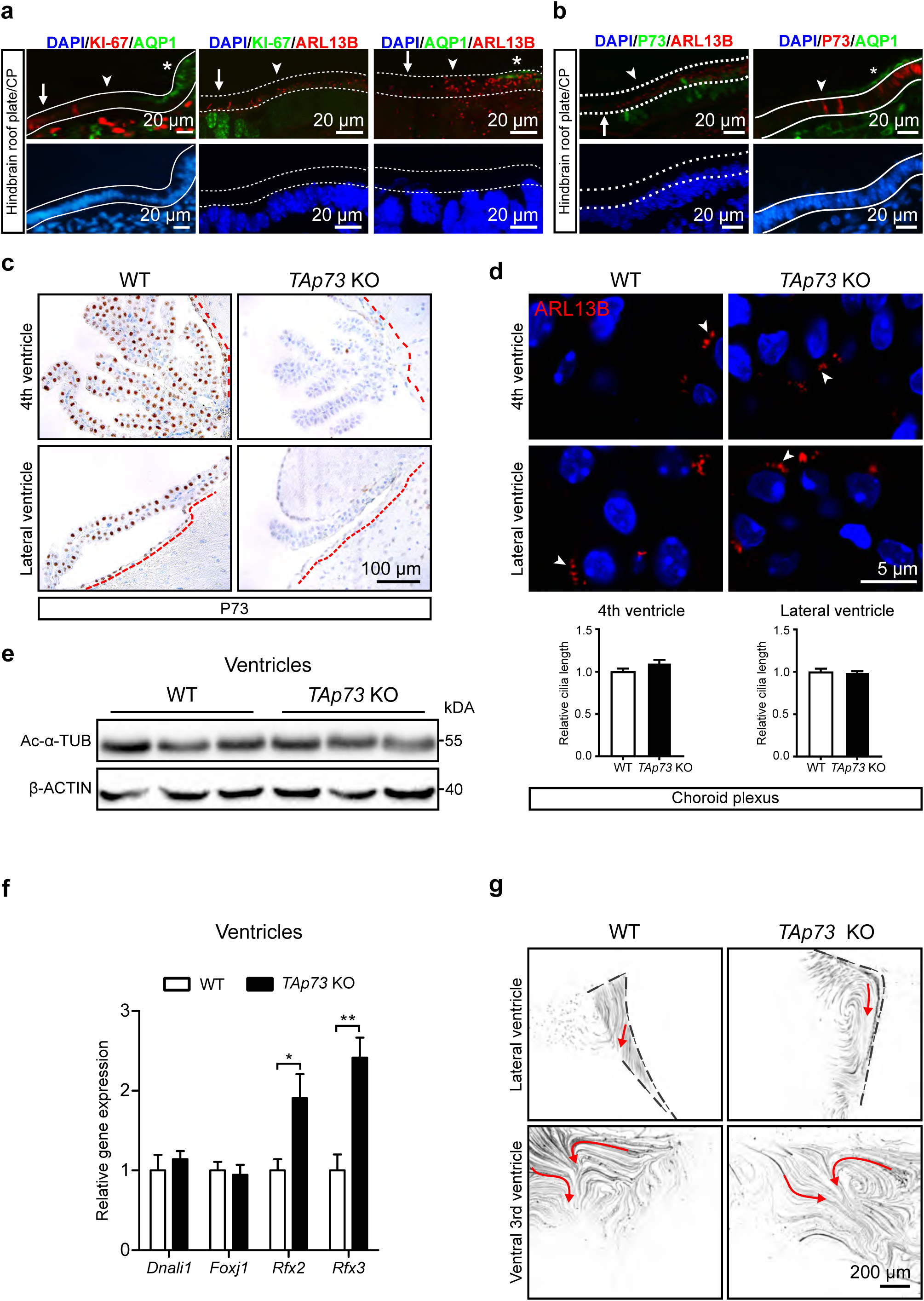
*TAp73* is dispensable for brain multiciliogenesis. (**a**) The expression of KI-67, Aquaporin 1 (AQP1, green), and ADP-ribosylation factor-like 13b (ARL13B, red) in WT hindbrain roof plate/CP at embryonic (E) day 14.5. Notice that KI-67^+^ roof plate progenitors, and AQP1^+^ CP epithelial cells are spatially separated. ARL13B labels monociliated roof pate progenitors and multiciliated CP epithelial cells. White lines demarcate roof pate epithelium (KI-67^+^/AQP1^-^, arrows), CP epithelium (KI-67^-^/AQP1^+^, asterisks), and “transition zone” (KI-67^-^/AQP1^-^, arrowheads) in which MCCs appear. Dotted lines mark apical cell surface with cilia. DAPI staining (blue) labels nuclei. (**b**) Expression of TAp73 (green, red), AQP1 (green), and ARL13B (red) in WT hindbrain roof plate/CP at E14.5. Dotted lines mark apical cell surface of roof plate (TAp73^-^, arrow) and transition zone (TAp73^+^, arrowhead). White lines mark transition zone (TAp73^+^/AQP1^-^, arrowhead) and CP epithelium (TAp73^+^/AQP1^+^, asterisk). DAPI staining (blue) labels nuclei. (**c**) Representative images of TAp73 expression of ependymal and CP epithelial cells in hindbrain and lateral ventricle from WT and *TAp73* KO. Red dotted lines mark ventricles lined with ependymal cells. Note that p73 expression is lost in *TAp73* KO mice. (**d**) Expression of the cilia marker ARL13B (red) in CP epithelial cells from WT and *TAp73* KO. White arrowheads mark cilia on cell surface. DAPI staining (blue) labels nuclei. Quantitation of average cilia length is shown in the lower panel. Data from a single experiment are shown (WT, *n*=12 cells [hindbrain] and 9 cells [lateral ventricle] from 2 mice; *TAp73* KO, *n*=17 cells [hindbrain] and 15 cells [lateral ventricle] from 3 mice). (**e**) Immunoblot analysis of Ac-α-TUB in brain ventricles from WT and *TAp73* KO animals. β-ACTIN serves as a loading control. Data are representative of two independent experiments. (**f**) Semi-quantitative PCR of *Dnali1, Foxj1, Rfx2*, and *Rfx3* in brain ventricles from WT and *TAp73* KO. Data from a single experiment are shown (WT, *n*=3; *TAp73* KO, *n*=4). (**g**) Movement of fluorescent beads along the ventricular system in WT and *TAp73* KO mice. Images of maximum intensity projections of representative movies of the lateral and the ventral 3^rd^ ventricles are shown (WT, *n*=2; *TAp73* KO, *n*=3; *TAp73* heterozygous, *n*=1). Red arrows mark the direction of bead flow. Bracket lines depict ependymal layer lining the lateral ventricles. Refer to **Supplementary Video S3a, b** for examples of recording of ciliary beating. All data are presented as mean ± SEM and relative to the WT group with \**P*<0.05, ** *P* <0.01.

The expression of TAp73 in ependymal and CP epithelial cells, along with recent studies demonstrating the role of E2F4/MCIDAS in multiciliogenesis of ependymal cells [7,48,49], led us to examine the role of TAp73 in MCCs in the brain. Immunostainings confirmed the loss of TAp73 expression in ependymal cells and the CP from *TAp73* KO mice (**Fig. 4c**), whereas morphological analysis revealed no apparent defect in these cells (**Supplementary Fig. 5a**). We performed immunostainings for the cilia markers ADP-ribosylation factor-like 13b (ARL13B) [50], Ac-α-TUB, and DNAI1 in the 4^th^ and lateral ventricles. In contrast to FTs and EDs, MCCs in ependyma and CP from *TAp73* KO animals are similar to those of WT mice (**Fig. 4d, e; Supplementary Fig. 5b-d**). No significant difference was observed in the expression of markers for epithelial differentiation of CP between control and *TAp73* KO animals (**Supplementary Fig. 6a-d**). RT-qPCR analysis demonstrated similar expression levels of *Dnali1* and *Foxj1*, whereas increased *Rfx2* and *Rfx3* mRNA levels were observed in brain ventricles from *TAp73* KO mice (**Fig. 4f**). Consistently, ciliary beating and bead flow in the cerebrospinal fluid appeared unaffected by *TAp73* loss (**Fig. 4g; Supplementary Video 3a, b**). Taken together, these results indicate that, unlike EDs, FTs, and the airways [12], the differentiation and function of MCCs in the brain remain intact despite *TAp73* loss.

### TAp73 regulates *miR-34/449* family members in diverse MCCs

Functional MCCs in the brain from *TAp73* KO mice suggest that other ciliogenic factors may rescue brain multiciliogenesis in the absence of *TAp73*. TAp73 influences post-transcriptional mechanisms *via* regulation of miRNAs [12]. Analysis of small RNA species from brain ventricles in *TAp73* KO mice revealed reduced *miR34bc* levels, along with a strong induction of the *miR449* cluster that works together with *miR34bc* to regulate multiciliogenesis in different tissues across species (**Fig. 5a, c; Supplementary Table 6**) [11,33,51–53]. In the brain, *miR449* is predominantly detected in the CP [54], where its expression undergoes >10 fold increase upon *TAp73* loss, whereas *miR34bc* levels strongly decline (**Fig. 5b, c**). Although *miR34b* levels were down-regulated as well in trachea from *TAp73* KO (**Supplementary Fig. 7a)**, *miR449* induction was less pronounced and more variable in FTs and EDs (**Fig. 5d**). Altogether, these results reveal a conserved reaction from the *miR-34/449* family following *TAp73* loss in diverse multiciliated epithelia.

**Fig. 5.**
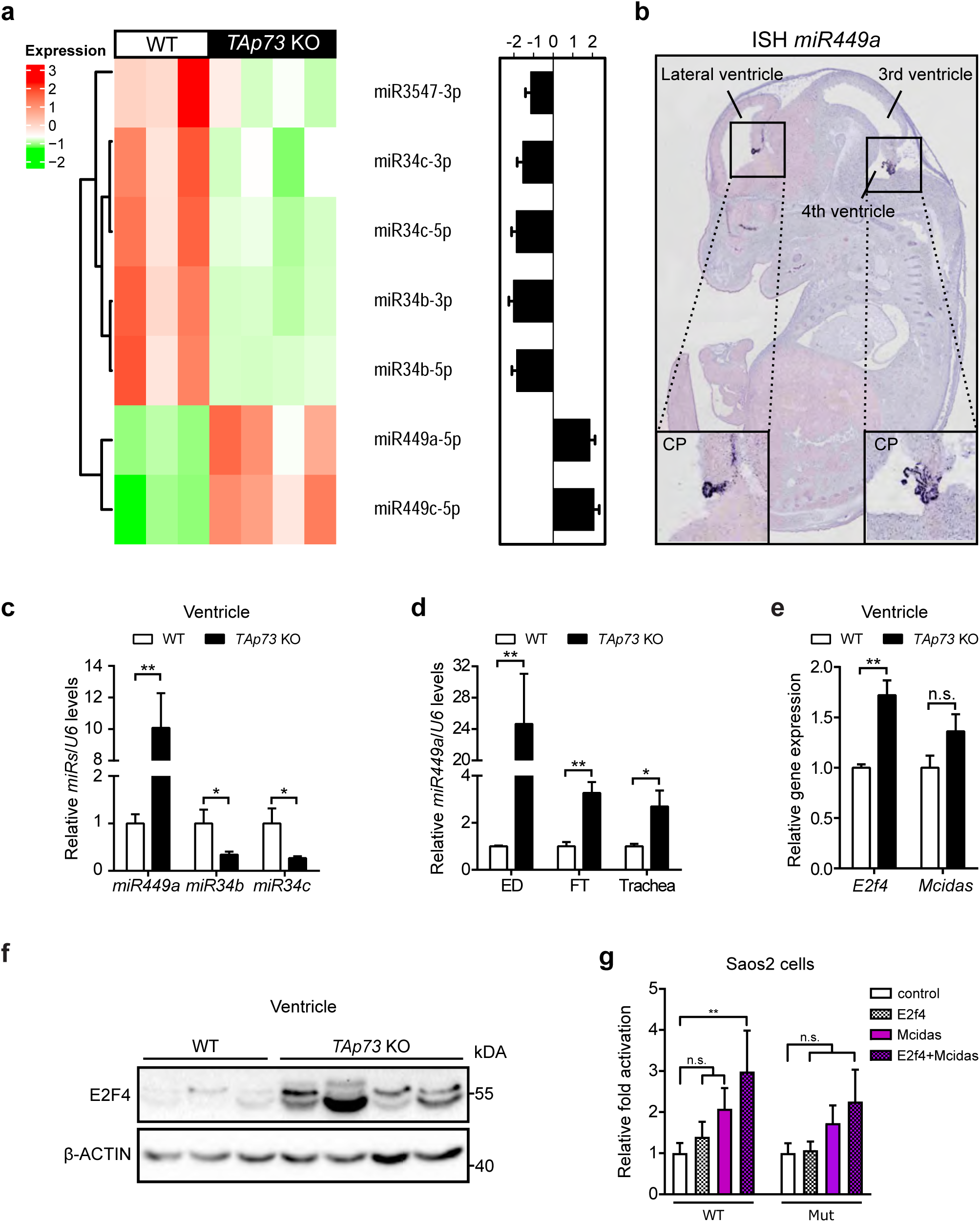
*TAp73* loss leads to changes in *miR-34/449* family and E2F4/MCIDAS circuit in the brain. (**a**) Hierarchical clustering of differentially expressed miRNAs in brain ventricles from WT and *TAp73* KO mice (WT, *n*=3; *TAp73* KO, *n*=4, one-way ANOVA, FDR < 0.05, fold change is shown). log2 values for miRNAs are plotted on the right. (**b**) *In situ* hybridization analysis of the expression of *miR449* in WT roof plate/CP at E14.5 (http://www.eurexpress.org/ee/) [77]). Semi-quantitative PCR analysis of *miR449a, miR34b*, and *miR34c* in brain ventricles (**c**), *miR449a* expression in EDs, FTs, and trachea (**d**), and *E2f4* and *Mcidas* levels in brain ventricles (**e**) from WT and *TAp73* KO mice. Data from a single experiment are shown (WT: ED, *n*=3; FT, *n*=7; trachea, *n*=4; ventricle, *n*=3. *TAp73* KO: ED, *n*=4; FT, *n*=8; trachea, *n*=4; ventricle, *n*=4). (**f**) Immunoblot of E2F4 in WT and *TAp73* KO ventricles. β-ACTIN serves as a loading control. Representative result of three independent experiments. (**g**) Luciferase assay of *miR449* regulatory regions containing E2F binding motifs. Three consensus E2F binding sites in *miR449* locus (http://jaspar.binf.ku.dk/) were placed in front of a luciferase cassette. A deletion mutant (Mut) that lacks the strongest consensus site was also created (**Supplementary Table 5**). WT or Mut luciferase vector was then co-transfected with empty vector (control), or vectors expressing E2F4, MCIDAS, or both. Fold changes in luciferase activities relative to those of control vector are shown. Date from 5 independent experiments are shown. All data are presented as mean ± SEM and relative to the WT group with \**P*<0.05, ** *P* <0.01.

In an effort to understand *miR449* upregulation we analyzed potential changes in the RB-E2F pathway known to regulate *miR449* levels [55,56]. However, expression of *E2f1, E2f3, Cdkn1a*, and *Cdkn1b* in brain ventricles were comparable between WT and mutant animals, (**Supplementary Fig. 8a, b**), indicative of a RB-E2F pathway unaffected by *TAp73* loss in brain MCCs. Interestingly, transcript and protein levels of the other E2F family member E2F4, which is a potent inducer of multiciliogenesis [6–8,48,57,58], were markedly increased in *TAp73* KO ventricles, despite only a mild increase of its cofactor *Mcidas* (**Fig. 5e, f**). In contrast, E2F4 levels in FTs and EDs were unaltered and even downregulated in tracheae (**Supplementary Fig. 7b-e**). Therefore, increased E2F4 levels concurrent with a *miR449* increase are restricted to the brain in *TAp73* KO mice.

To assess potential E2F4 contribution to *miR449* elevation, we used the genomic region of *miR449* containing three putative E2F binding sites in a reporter-based assay. Indeed, E2F4 in combination with MCIDAS elicited a strong transcriptional response from the *miR449* locus, a reaction almost abolished by mutating the strongest out of three E2F consensus motifs (**Fig. 5g; Supplementary Table 5**). Together, these results indicate that increased E2F4/MCIDAS activity may stimulate *miR449* expression in *TAp73* KO brains.

### TAp73 collaborates with *miR449* in brain multiciliogenesis

Our data suggest that *miR449* upregulation may compensate at least partially for *TAp73* loss to maintain brain multiciliogenesis. To address this, we generated mice with a deletion of the *miR449* cluster in addition to *TAp73*. Strikingly, *TAp73*^*-/-*^;*miR449*^*-/-*^ (*TAp73xmiR449* KO) mice developed severe hydrocephalus (**Fig. 6a; Supplementary Fig. 9a**). Since defective ependymal and CP cilia contribute to the development of hydrocephalus [59–61], we next assessed ciliation in the ventricles of *TAp73*x*miR449* KO mice. Analysis of the expression of ARL13B in CP epithelium revealed a decrease in cilia number and length in the absence of *miR449*, whereas a more pronounced reduction in cilia was observed in *TAp73xmiR449* KO mice (**Fig. 6b, c; Supplementary Fig. 9b**). TEM studies also revealed mildly disorganized apical docking of basal bodies in ependymal cells in *TAp73* KO and *TAp73*x*miR449* KO mice (**Fig. 6d; Supplementary Fig. 9c**); however, Ac-α-TUB content was similar in ependymal cells among WT and *TAp73*x*miR449* KO animals (**Supplementary Fig. 9d**). Consistently, ciliary beating and bead flow over ventricles appeared unaffected in *TAp73*x*miR449* KO animals (**Supplementary Fig. 9e; Supplementary Video 3a, c**). Furthermore, expression of cytokeratins, AQP1, and OTX2 in CP epithelial cells was similar among WT, *miR449* KO, and *TAp73xmiR449* KO animals (**Supplementary Fig. 10a-c**). Despite the role of Notch signaling in CP development and tumorigenesis [31,62], RNAscope studies revealed similar expression of NOTCH targets *Hes1* and *Hes5* in the roof plate of WT, *miR449* KO, and *TAp73xmiR449* KO embryos at day E14.5 (**Supplementary Fig. 11)**. In summary, additional loss of *miR449* in *TAp73* KO mice strongly impairs ciliogenesis in the CP, but only slightly affects ependymal cilia, which is consistent with its prominent expression in the CP [54] (**Fig. 5b**). Thus, our data indicate that *miR449* collaborates with *TAp73* to drive multiciliogenesis in the brain.

**Fig. 6.**
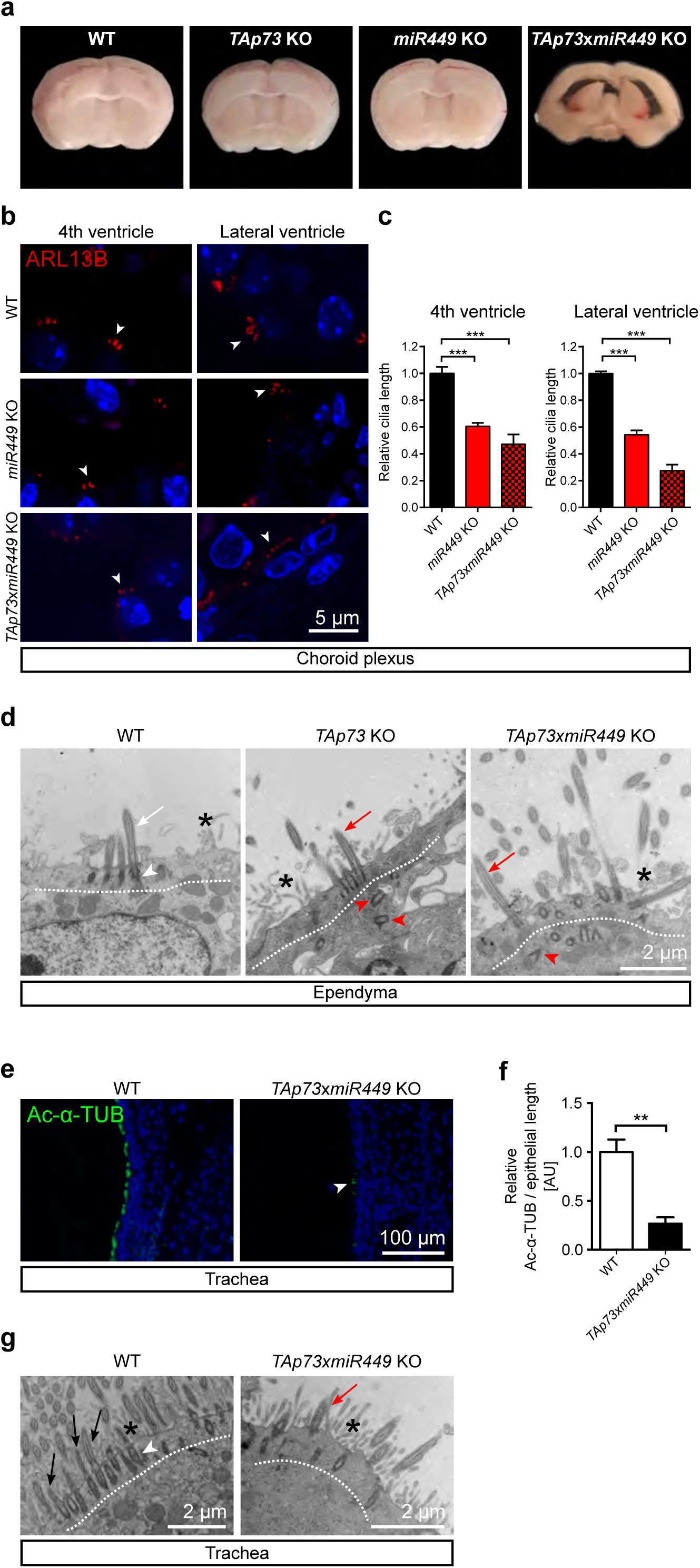
TAp73 functions through *miR449* in brain multiciliogenesis. (**a**) Coronal brain slices from WT, *TAp73* KO, *miR449* KO and *TAp73xmiR449* KO mice. Note that *TAp73xmiR449* KO mice display enlarged lateral ventricles. (**b**) ARL13B (red) expression in CP epithelial cells of the 4^th^ and lateral ventricles from WT, *miR449* KO, and *TAp73xmiR449* KO animals. White arrowheads mark cilia on cell surface. DAPI staining (blue) labels nuclei. (**c**) Quantitation of average cilia length of CP epithelial cells shown in (**b**). Data from a single experiment are shown (WT, *n*=4 cells [hindbrain, lateral ventricle] from 2 mice; *miR449* KO, *n*=18 cells [hindbrain] and 14 cells [lateral ventricle] from 4 mice; *TAp73xmiR449* KO, *n*=8 cells [hindbrain, lateral ventricle] from 3 mice). (**d**) Representative TEM photomicrographs of ependymal cells in WT, *TAp73* KO, and *TAp73*x*miR449* KO mice. Dotted lines mark apical region of the cells. Notice that WT cells possess cilia (white arrow) and basal bodies (white arrowhead) docked to the apical surface, whereas mutant cells have a similar number of cilia (red arrows) but disorganized basal bodies (red arrowheads) located further away from the apical surface. Interspersed microvilli are marked with asterisks. (**e**) Representative staining of Ac-α-TUB (green) in tracheae from WT and *TAp73xmiR449* KO mice. DAPI staining (blue) labels nuclei. Note that mutants harbor less and shorter cilia (arrowhead) than WTs. (**f**) Quantitation of Ac-α-TUB signals normalized to epithelial length is shown. Data from a single experiment are shown (*n*=4 samples/genotype). (**g**) Representative TEM photomicrographs of tracheae from WT and *TAp73xmiR449* KO mice. Dotted lines mark apical region of the cells. Notice the abundant cilia (black arrows) and clustered basal bodies (white arrowhead) docked to apical surface in WT cells, whereas mutant cells exhibit fewer cilia (red arrow). Interspersed microvilli are marked with asterisks. All data are presented as mean ± SEM and relative to the WT group with ** *P* <0.01, *** *P* <0.001.

**Fig. 7.**
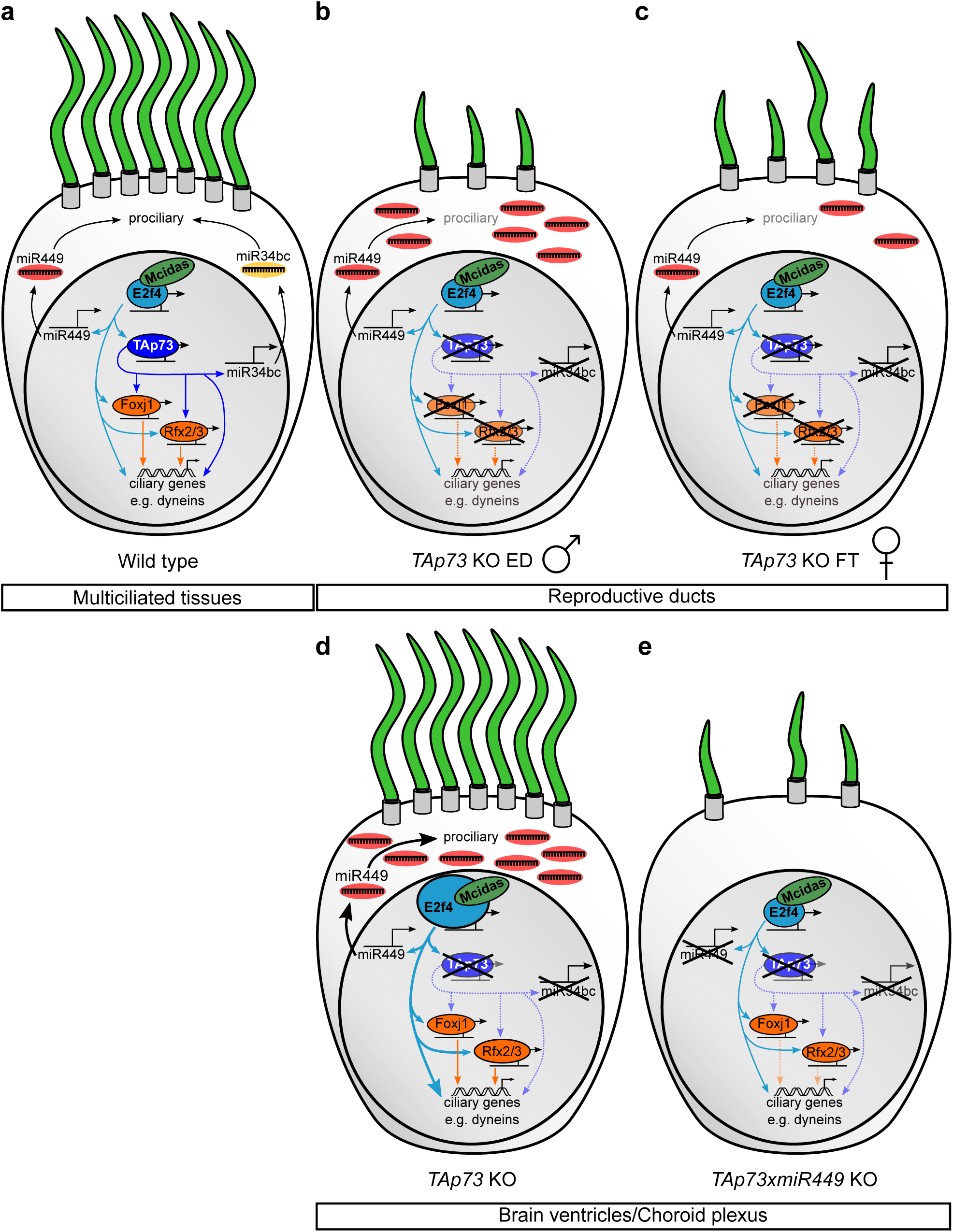
Schematic diagram of the molecular circuits of TAp73-driven multiciliogenesis in diverse tissues. (**a**) TAp73-dependent transcriptional network, including dyneins, *miR34bc, Foxj1, Rfx2, and Rfx3* factors, critically regulates multiciliogenesis in various ciliated epithelia downstream of *E2f4*/*Mcidas*. In the EDs (**b**) and FTs (**c**) *TAp73* KO impairs multiciliogenesis concurrent with male and female fertility. (**d**) TAp73 is not essential for multiciliogenesis in the brain; however, *TAp73* loss leads to upregulation of pro-ciliogenic *E2f4* and its target *miR449*. (**e**) Further removal of *miR449* in *TAp73* KO animals leads to reduced number and length of CP cilia and severe hydrocephalus, indicating that *miR449* and TAp73 complement each other to support brain ciliogenesis.

As *miR449* was induced upon *TAp73* deletion in further multiciliated tissues, we analyzed tracheae and EDs in *TAp73xmiR449* KO mice. Immunostainings and TEM consistently revealed a dramatic decrease in cilia coverage and an increase in defective basal body docking in trachea from *TAp73xmiR449* KO animals compared to WT animals (**Fig. 6e-g; Supplementary Fig. 12a**), a phenotype bearing resemblance to our previous findings in the airways of *TAp73* KO animals [12]. Likewise, loss of *miR449* did not further enhance MCC reduction in *TAp73*-deficient EDs (**Supplementary Fig. 12b, c**). Thus, additional deletion of the *miR449* cluster fails to exacerbate ciliary defects in trachea and EDs in the absence of *TAp73*.

Overall, our data indicate that TAp73 utilizes the unique topology of its transcriptional circuit to communicate with the *miR-34/449* family and other crucial regulators of motile multiciliogenesis e.g. *E2F4/MCIDAS* to regulate brain multiciliogenesis (**Fig. 7a, d, e**).

## Discussion

TAp73 activates a plethora of ciliogenic effectors to drive multiciliogenesis in the airways [12,13]. The current study examines the role of TAp73-driven molecular circuit in MCCs of reproductive tracts and the brain. Our results revealed a profound reduction of cilia in EDs and FTs from *TAp73*-deficient mice, as well as diminished *Foxj1, Rfx2, and Rfx3* expression. These molecular and cellular changes in MCCs are reminiscent of our previous findings in respiratory epithelia of these mice, suggesting that male and female infertility associated with *TAp73* loss could be in part related to the observed cilia loss. The expression of the axonemal dyneins *Dnai1* and *Dnali1*, both of which exhibit TAp73 binding in their genomic loci, was also significantly reduced in EDs and FTs from mutant animals, indicating that they are part of the TAp73-directed multiciliogenesis program in reproductive tracts.

Consistent with previous reports, we found partial degradation of the germinal epithelium and reduced sperm cell production in *TAp73* KO mice [42,43]. The EDs are comprised of MCCs, which are required for fluid circulation and reabsorption, thereby facilitating the transport of spermatozoa to their storage and maturation in the epididymis [22,24,25]. Despite the presence of flagellated spermatozoa in testis, lack of spermatozoa in epididymis of *TAp73* KO mice indicates that defective multiciliogenesis may contribute to male sterility. Indeed, disruption of transcriptional regulators of multiciliogenesis has been shown to cause infertility in mice and humans [3,63], whereas fertility issues have been reported in female primary ciliary dyskinesia patients [20,21]. Importantly, *TAp73* is downregulated as women age [64], and certain single nucleotide polymorphisms in *TP73* are associated with female patients over 35 years of age seeking *in vitro* fertilization [65,66]. Hence, the integrity of MCCs is critical for reproductive health. Further studies using tissue-specific deletion of *TAp73* in MCCs of EDs and oviducts are necessary to delineate its role in reproductive motile cilia maintenance and fertility.

In the brain, *TAp73* expression is initiated at the onset of multiciliated differentiation of ependymal and CP epithelial cells. However, our data indicate that *TAp73* is dispensable for the generation of cilia in the brain, although it is plausible that *TAp73* loss results in more subtle defects such as polarity and cilia orientation [67,68]. In contrast to the dynamic TAp73-dependent program in the airways and reproductive tracts, expression of *Foxj1, Rfx2, and Rfx3* in the brain remains mostly unaltered in the absence of *TAp73*, suggesting that other effectors maintain the activity of the molecular circuit to support MCC differentiation.

Previous studies revealed robust expression of *GemC1* and *E2f/Mcidas*, all of which are capable of transcriptional activation of *Foxj1, TAp73* itself, and many other ciliogenic effectors e.g. *Rfx2* and *Rfx3* in MCCs of the brain [4,6,8,48,69]. Indeed, E2F4/MCIDAS expression is upregulated in the brain but not in other multiciliated tissues upon *TAp73* loss, and therefore may facilitate brain multiciliogenesis. In agreement, loss of either *Mcidas* or *GemC1*, both transcriptional activators of TAp73, leads to defect in MCC differentiation and hydrocephalus [3,6].

Although it is less clear how *TAp73* loss results in enhanced E2F/MCIDAS activity in the brain, a quick look downstream of TAp73 provides some clues: reduced expression of the TAp73 target *miR34bc* is concurrent with an induction of *miR449* in the absence of *TAp73*. Interestingly, expression of Cdkn1a/p21, Cdkn1b/p27, E2f1, and E2f3 in brain ventricles remain unchanged following *TAp73* loss, suggesting that *TAp73* loss regulates E2F and *miR449* activity independently of the conserved RB-E2F1 axis. *miR449* induction is commonly observed in *miR34*-deficient MCCs, whereas ablation of the entire *miR-34/449* family severely impairs multiciliogenesis in diverse tissues [33,70]. *miR449* is known to inhibit the NOTCH pathway to relieve the suppression of MCC fate determination; however, NOTCH pathway activity in the CP remains unchanged after *miR449* loss. Given the diverse targets of the *miR-34/449* family, it is plausible that *miR449* may indirectly increase E2F/MCIDAS activity in MCCs of the brain independent of NOTCH inhibition. Conversely, transcriptional activation of *miR449* by E2F/MCIDAS complexes may complete the feedback loop to keep the molecular circuit fully engaged in the absence of *TAp73*.

This interpretation posits that the crosstalk between *miR449* and E2F/MCIDAS serves as a crucial backup circuit for TAp73-driven multiciliogenesis network in the brain. Indeed, combined deletion of *TAp73* and *miR449* results in disruption of multiciliogenesis in the brain and hydrocephalus, defects distinct from those associated with complete loss of the *miR-34/449* family [33,71]. In *TAp73*-deficient MCCs outside the brain that exhibit less prominent increase in *miR449* and no increase in E2F4 levels, further deletion of *miR449* cluster fails to exacerbate multiciliogenesis defects caused by *TAp73* loss, indicating that TAp73 functions at least partially through *miR449* to support MCCs in the brain. Recent studies also demonstrated the role of *TAp73*-driven *miR34a* expression in neuronal development [72]. Therefore, interaction of TAp73 with *miR-34/449* family members is crucial for normal brain functions.

Nonetheless, detailed studies are necessary to clarify the interaction between *miR449* and E2F4/MCIDAS pathway in MCCs in the brain, but also to address *miR449* regulation in *TAp73*-deficient MCCs outside the brain.

Unlike *TAp73* mutant animals, *p73* KO mice lacking both *TAp73* and *ΔNp73* exhibit hydrocephalus, defective ependymal cell maturation and aqueduct stenosis, suggesting a potential role for *ΔNp73* in ependymal cells [73,74]. Given the abnormal apical localization of basal bodies in ependymal cells along with ciliary defects in the CP from *TAp73xmiR449* KO mice, it is conceivable that *ΔNp73* may regulate *miR449* expression indirectly in these cells. In support of this notion, *miR449* is highly expressed in the CP whereby its loss alone leads to ciliary defects, whereas *ΔNp73* deletion also results in defects in the CP [75]. Further analysis of the multiciliogenesis network and *miR449* expression in MCCs in the brains of *p73* KO and *ΔNp73* KO animals are necessary to resolve these questions.

## Acknowledgments

We thank Tak Mak for providing *TAp73* KO mice, Gerd Hasenfuß for support, Matthias Dobbelstein for hosting and Karola Metze and Verena Siol for assistance. M.L. is supported by Deutsche Forschungsgemeinschaft (DFG Li 2405); H.Z. by New York Institute of Technology, Sanford Research, Matthew Larson Foundation, Institutional Development Award from the National Institute of General Medical Sciences under grant numbers 5P20GM103548, 1P20GM103620-01A1, and National Cancer Institute (R01CA220551); F.B. by Wilhelm-Sander-Stiftung (2016.041.1); A.K.G by the Max Planck Society. We thank Heymut Omran’s group for introduction to cilia microscopy and Travis Stracker for disclosure of non-published data.

## Author Contributions

Me.W. and T.E. characterized cilia defects and gene expression and generated figures. Me.W. and Ma.W. validated TAp73 targets by WB and ChIP. E.E. and F.B. contributed IF analysis of human epididymis. E.E. performed cilia quantification on tracheae. D.R. performed electron microscopy analysis. C.W. maintained mice, performed RNA isolation and qPCRs. L.V-H. and S.B. contributed to Western blot analysis of different tissues. Z. H. performed RNAscope analysis for *TAp73* on diverse tissues. K.B.G., J.Z., L.L., and H.Z. contributed brain analyses. A-K.G analyzed *ex vivo* ciliary beating. O.S. analyzed small RNA sequencing data. S.A. contributed to interpretation and supported the group. M.L. developed the project, interpreted the data, designed and coordinated the experiments to complete this study. Me.W., T.E., H.Z., and M.L. were major contributors to manuscript preparation.

## Electronic supplementary material

The online version of this article contains supplementary material, which is available to authorized users.

## Supplementary information

### Supplementary Figures

**Supplementary Fig. 1.**
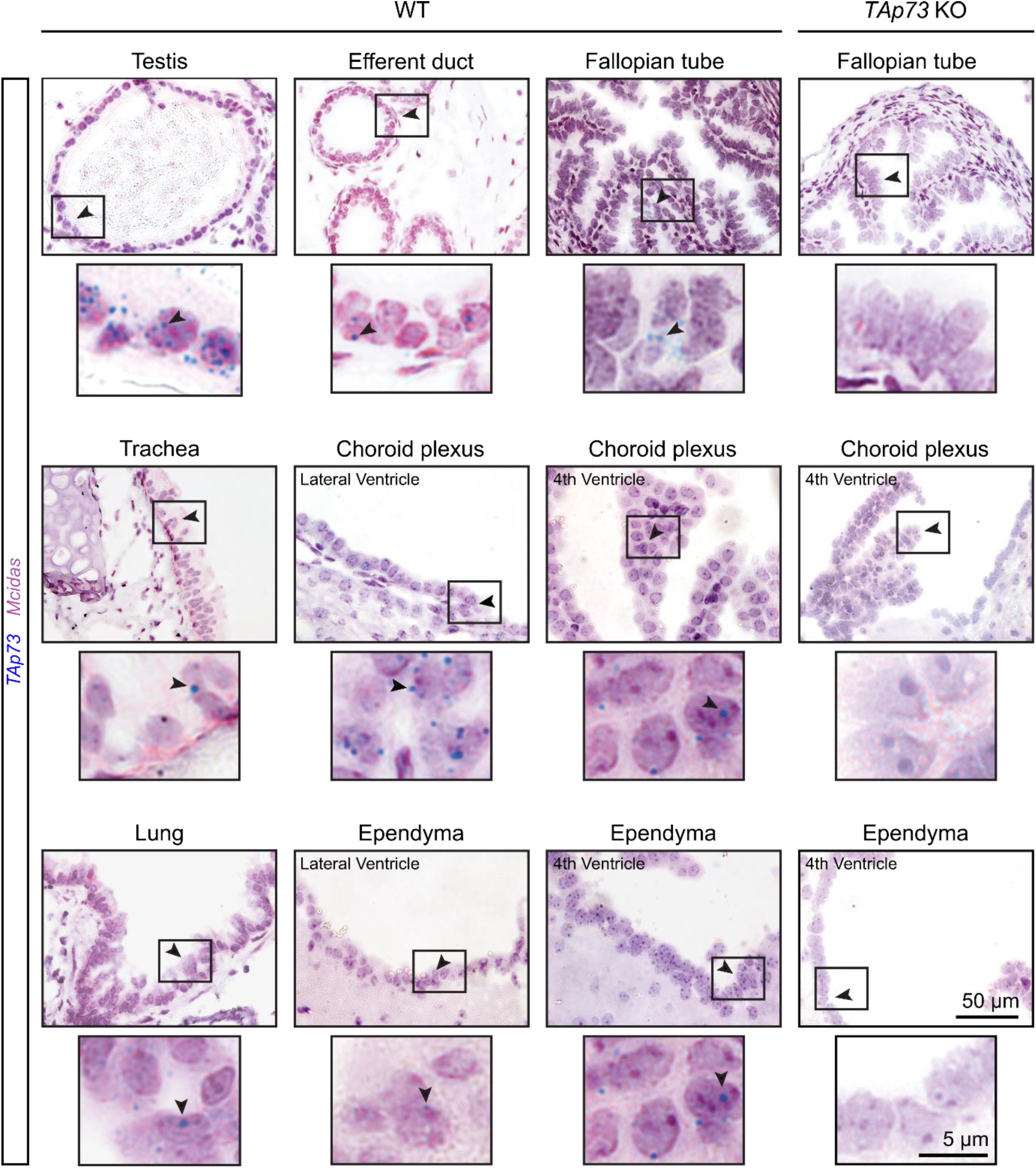
TAp73 is expressed in diverse multiciliated tissues. RNAscope analysis of *TAp73* (blue) and *Mcidas* (red) expression in testis, efferent ducts, fallopian tube, trachea, lung, ependymal and choroid plexus epithelial cells in hindbrain and lateral ventricle from adult wild type (WT) and *TAp73* knockout (KO) mice (3 months of age). Arrowheads mark cells in boxed regions that are also shown in higher magnification. Notice that *TAp73* is absent in *TAp73* KO tissues.

**Supplementary Fig. 2.**
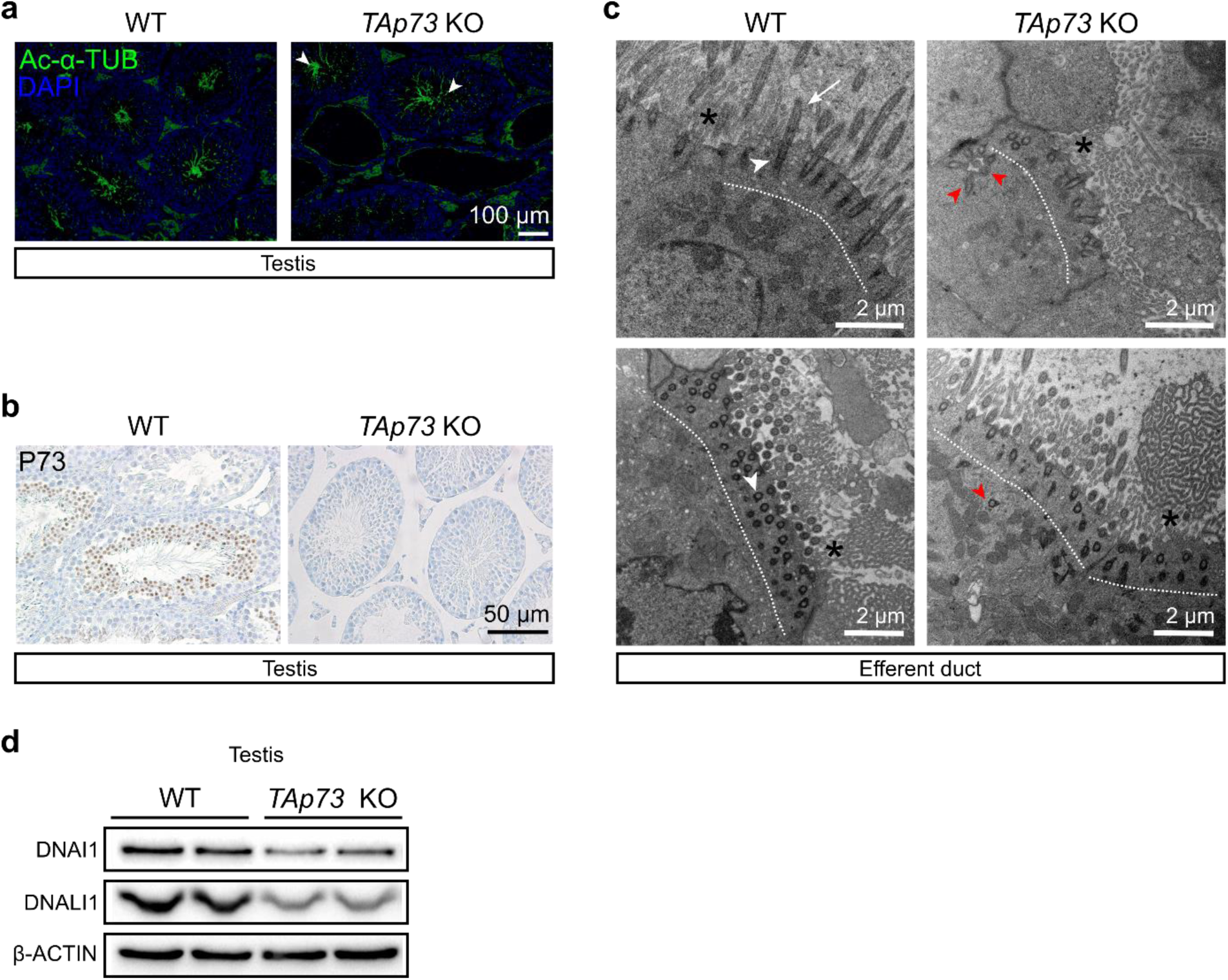
Loss of *TAp73* impairs multiciliogenesis in the male reproductive duct. (**a**) Representative images of the expression of acetylated alpha-tubulin (Ac-α-TUB, green) in testes from WT and *TAp73* KO mice. DAPI staining (blue) labels nuclei. Notice, although at a reduced level, spermatozoa (Ac-α-TUB^+^) are present in mutant testis (arrowheads). (**b**) Expression of TAp73 in testes from WT and *TAp73* KO mice at 3 months of age, respectively. Notice that TAp73 is expressed in testes from WT mice, but absent in mutant testes. (**c**) Representative photomicrographs of transmission electron microscopy (TEM) of efferent ducts from WT and *TAp73* KO mice. Dotted lines mark apical region of the cells. Notice the abundant cilia (white arrow) and clustered basal bodies (white arrowheads) on WT cells, whereas mutant cells exhibit disorganized basal bodies (red arrowheads) located away from apical surface. Interspersed microvilli are marked with asterisks. (**d**) Immunoblot analysis of the expression of DNAI1 and DNALI1 in testes from WT and *TAp73* KO animals. β-ACTIN serves as a loading control. DNAI1 and DNALI1 levels are reduced in mutant testis compared to WT animals. Representative results of three independent experiments are shown.

**Supplementary Fig. 3.**
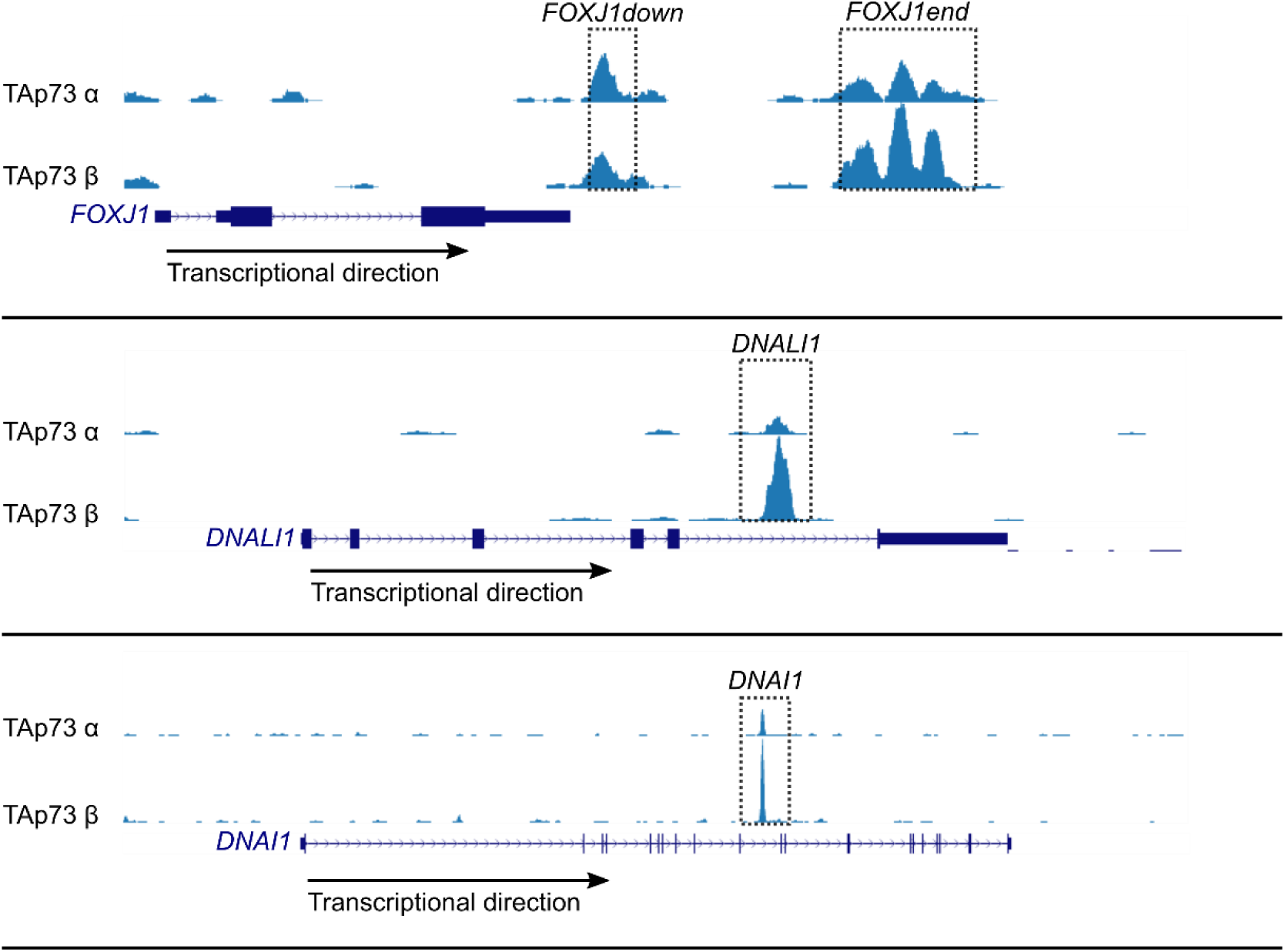
TAp73 is associated with ciliary genes. TAp73 binding at *FOXJ1, DNALI1*, and *DNAI1* genomic loci is shown in results from ChIP-seq [1], Geo accession no. **GSE15780**). Boxed regions mark genomic loci enriched with TAp73 binding and validated by ChIP-qPCR (**Fig. 2e**).

**Supplementary Fig. 4.**
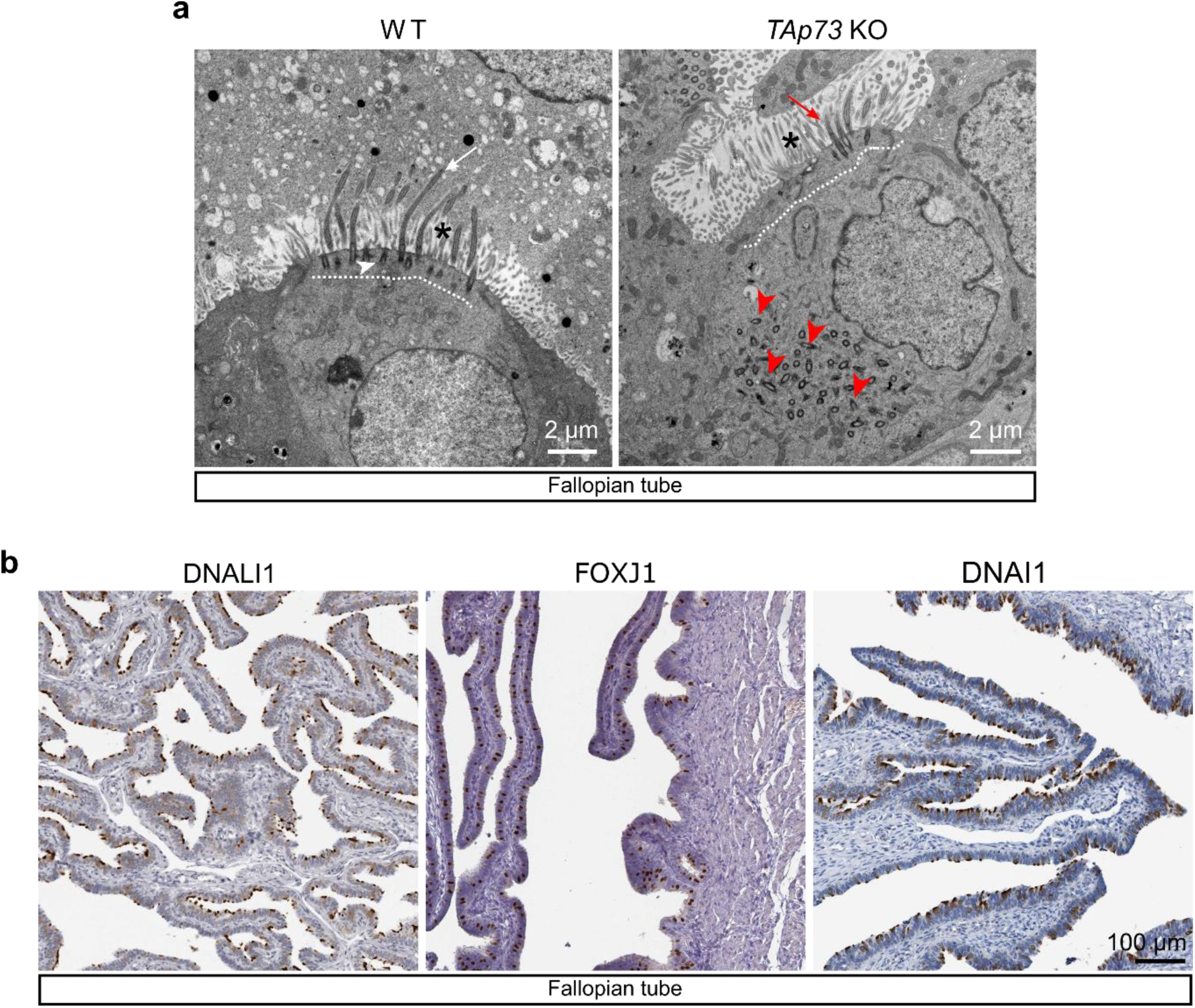
TAp73 KO mice show defective motile cilia in the fallopian tube. (a) Representative photomicrographs of transmission electron microscopy (TEM) of fallopian tubes from WT and *TAp73* KO mice. Dotted lines mark apical region of the cells. Notice the presence of abundant cilia (white arrow) and clustered basal bodies (white arrowhead) docked to apical surface of the WT cell, whereas the mutant cell displays fewer cilia (red arrow) and disorganized basal bodies (red arrowheads) located away from apical surface. Interspersed microvilli are marked with asterisks. (**b**) Expression of DNALI1, FOXJ1, and DNAI1 in human FT epithelia. Images were retrieved from the Human Protein Atlas (DNALI1: http://www.proteinatlas.org/ENSG00000163879-DNALI1/tissue/fallopian+tube, FOXJ1: http://www.proteinatlas.org/ENSG00000129654-FOXJ1/tissue/fallopian+tube, DNAI1: http://www.proteinatlas.org/ENSG00000122735-DNAI1/tissue/fallopian+tube).

**Supplementary Fig. 5.**
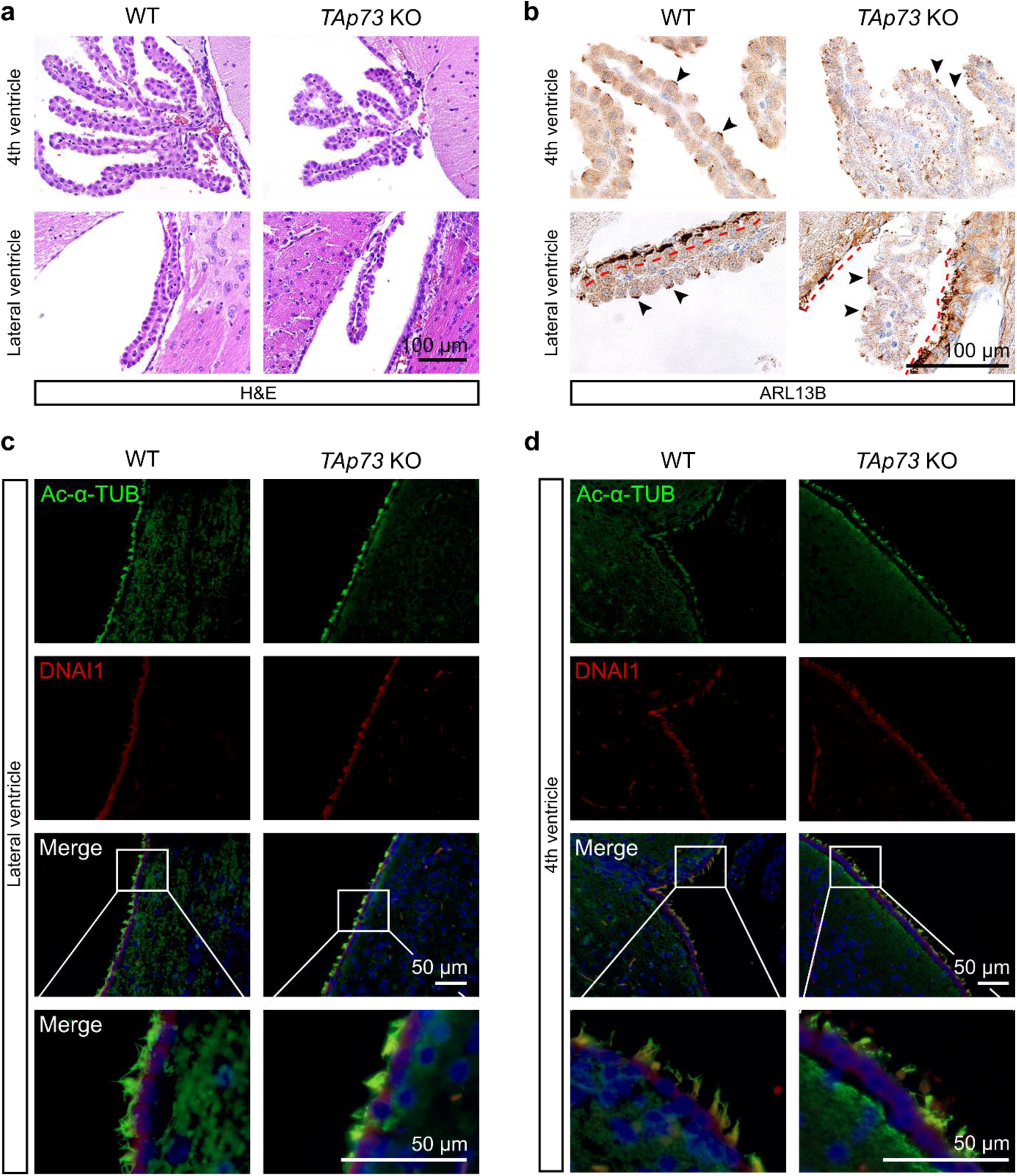
*TAp73* loss does not affect multiciliogenesis in the brain. (**a**) Representative images of H&E staining of choroid plexus (CP) in the 4^th^ and lateral ventricle from WT and *TAp73* KO animals. (**b**) Representative images of the expression of ARL13B in CP of the 4^th^ and lateral ventricle from WT and *TAp73* KO animals. Arrowheads mark cilia on CP epithelial cells. Red dotted lines delineate the boundary of lateral ventricles lined with ependymal cells. Expression of Ac-α-TUB (green) and DNAI1 (red) in ependymal cells of lateral ventricle (**c**) and the 4^th^ ventricle (**d**) from WT and *TAp73* KO animals. Boxed regions are magnified in bottom panels. DAPI staining (blue) labels nuclei.

**Supplementary Fig. 6.**
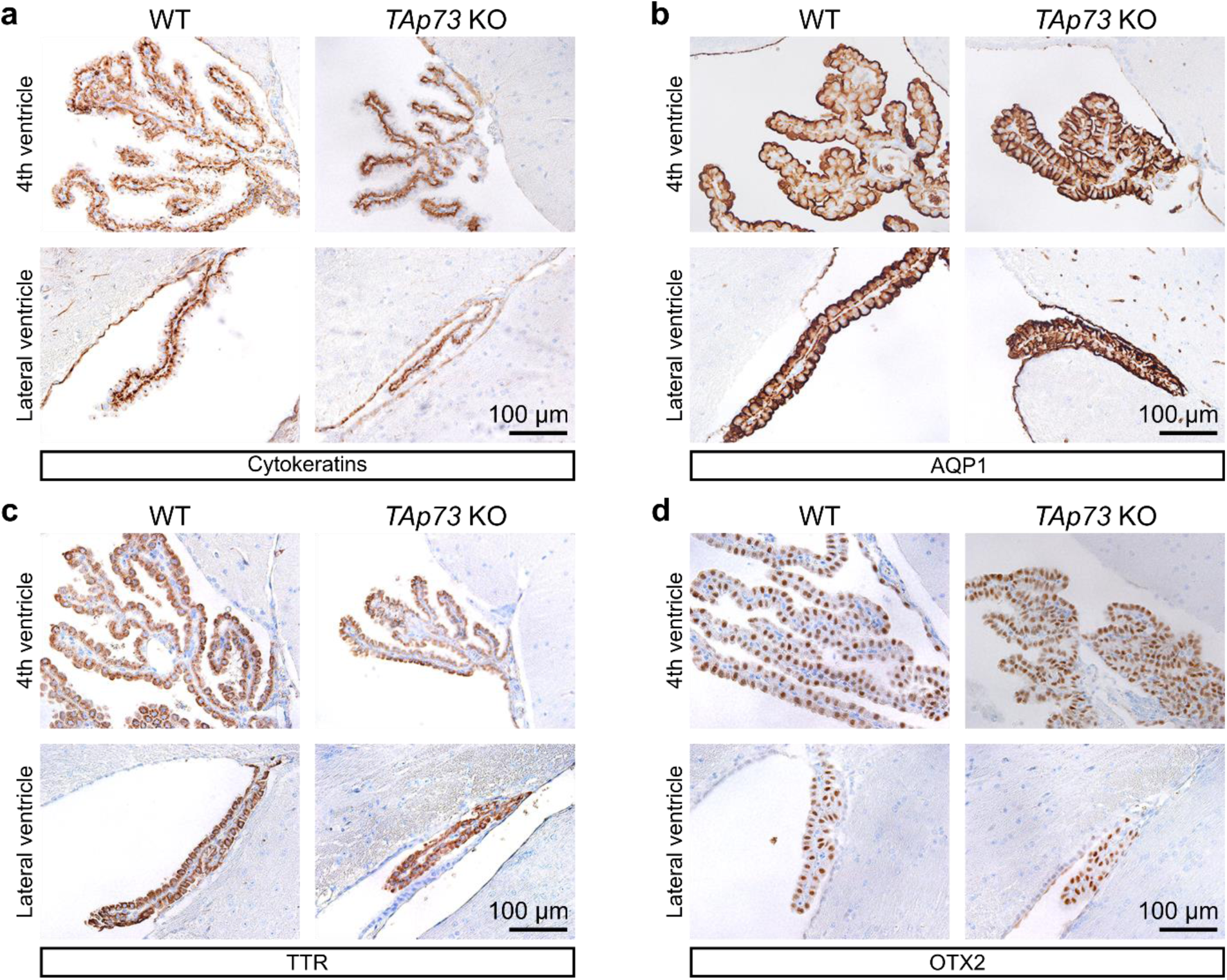
*TAp73* loss does not affect epithelial differentiation of choroid plexus cells. Representative images of the expression of cytokeratins (**a**), aquaporin 1 (AQP1, **b**), transthyretin (TTR, **c**), and orthodenticle homeobox 2 (OTX2, **d**) in CP epithelium of the 4^th^ and lateral ventricles from WT and *TAp73* KO animals.

**Supplementary Fig. 7.**
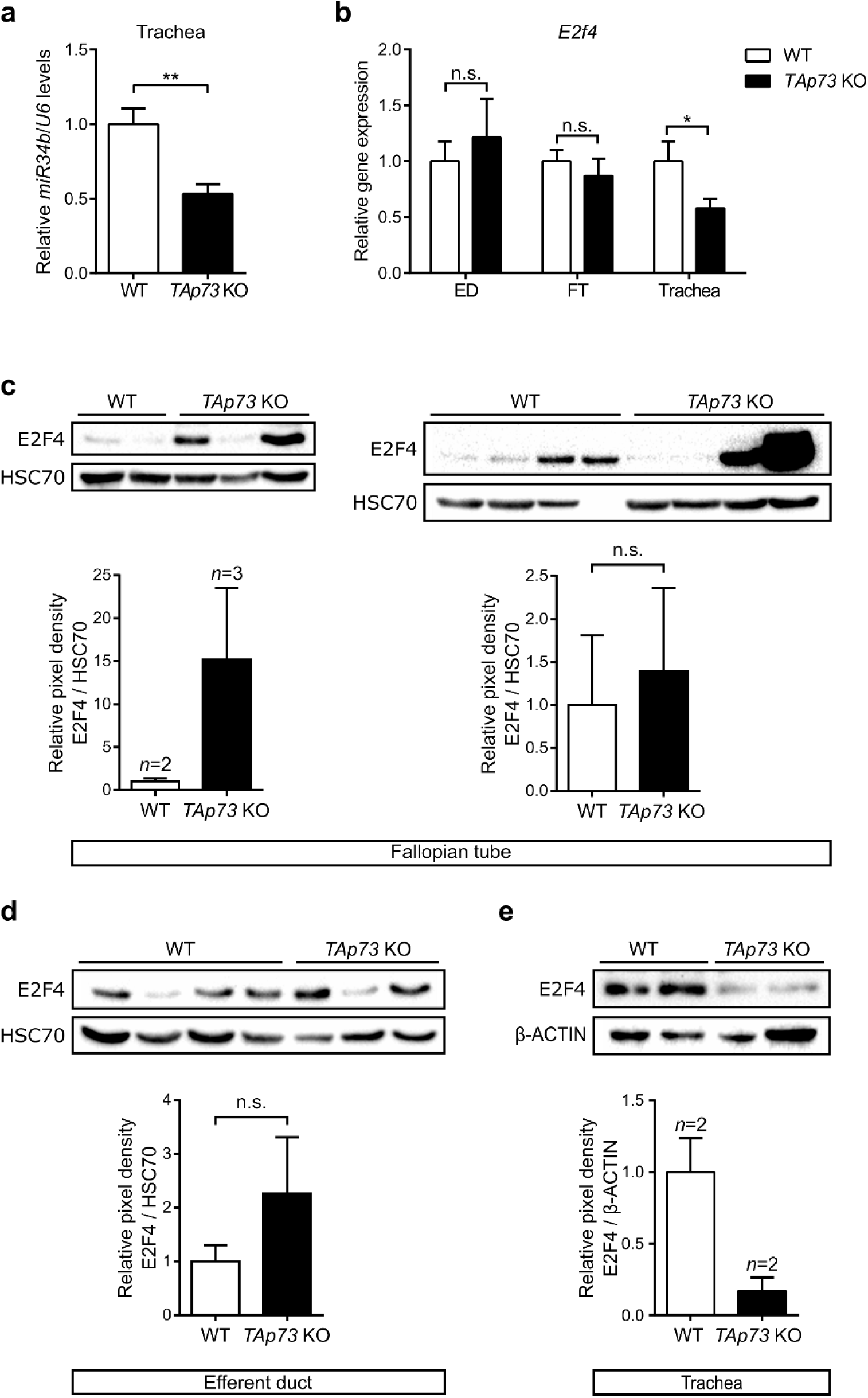
Analysis of gene expression in multiciliated tissues of *TAp73* KO animals. (**a**) Semi-quantitative PCR analysis of *miR34b* expression in trachea from WT and *TAp73* KO mice. Data from a single experiment are shown (WT: *n*=3; *TAp73* KO: *n*=4). (b) Semi-quantitative PCR analysis of *E2f4* expression in brain ventricles, efferent ducts (ED), fallopian tubes (FT), and tracheae from WT and *TAp73* KO mice. Data from a single experiment are shown (WT: ED, *n*=5; FT, *n*=7; trachea, *n*=4; *TAp73* KO: ED, *n*=4; FT, *n*=8; trachea, *n*=4). Immunoblot analysis of the expression of E2F4 in FTs (**c**), EDs (**d**), and tracheae (**e**) from WT and *TAp73* KO animals. β-ACTIN or HSC70 serve as loading controls. Quantitation of the signal intensity of E2F4 bands normalized to that of the loading control is shown below each immunoblot (n.s. non-significant). Representative results from three independent experiments are shown. All data are presented as mean ± SEM and relative to the WT group with \**P*<0.05, ** *P* <0.01.

**Supplementary Fig. 8.**
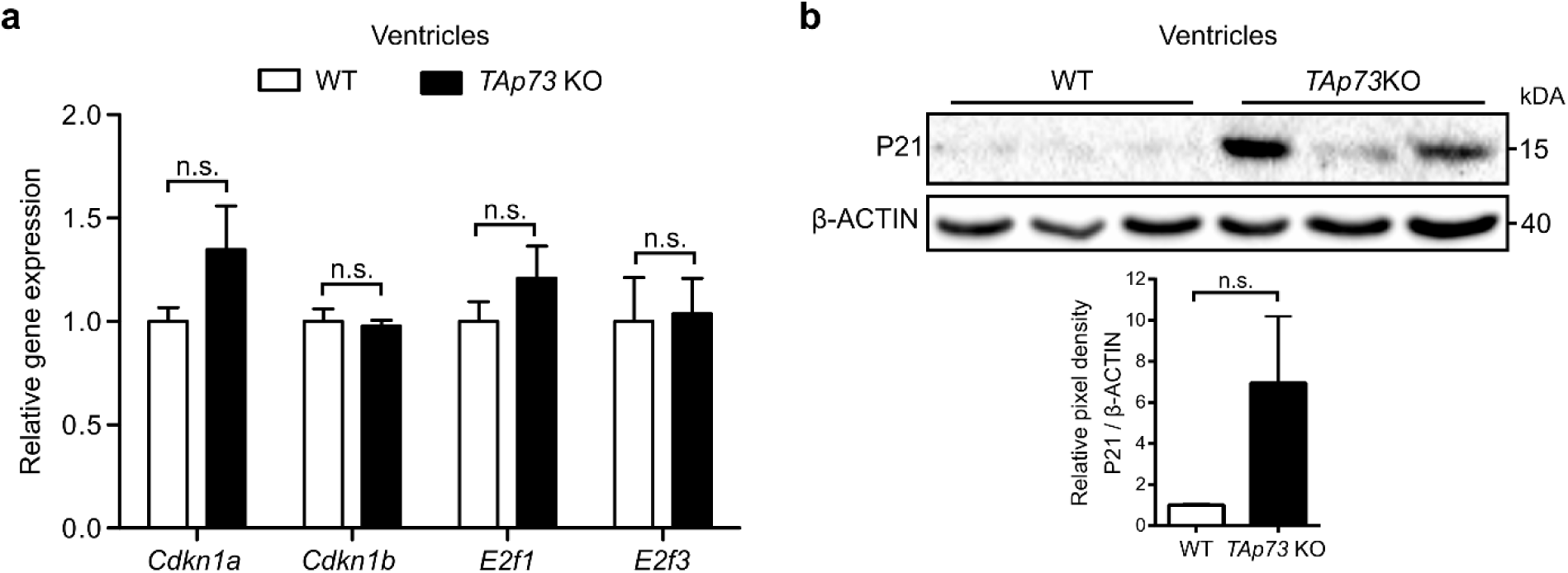
pRb/E2F pathway activity is not deregulated in *TAp73* KO ventricles. (**a**) Semi-quantitative PCR analysis of *Cdkn1a, Cdkn1b, E2f1*, and *E2f3* in brain ventricles from WT and *TAp73* KO animals. Data from a single experiment are shown (*n*=4). (b) Immunoblot analysis of P21 expression in brain ventricles from WT and *TAp73* KO animals. Data are representative of three independent experiments. Quantitation of the signal intensity of P21 bands relative to that of β-ACTIN is shown (*n*=3). All data are presented as mean ± SEM and relative to the WT group.

**Supplementary Fig. 9.**
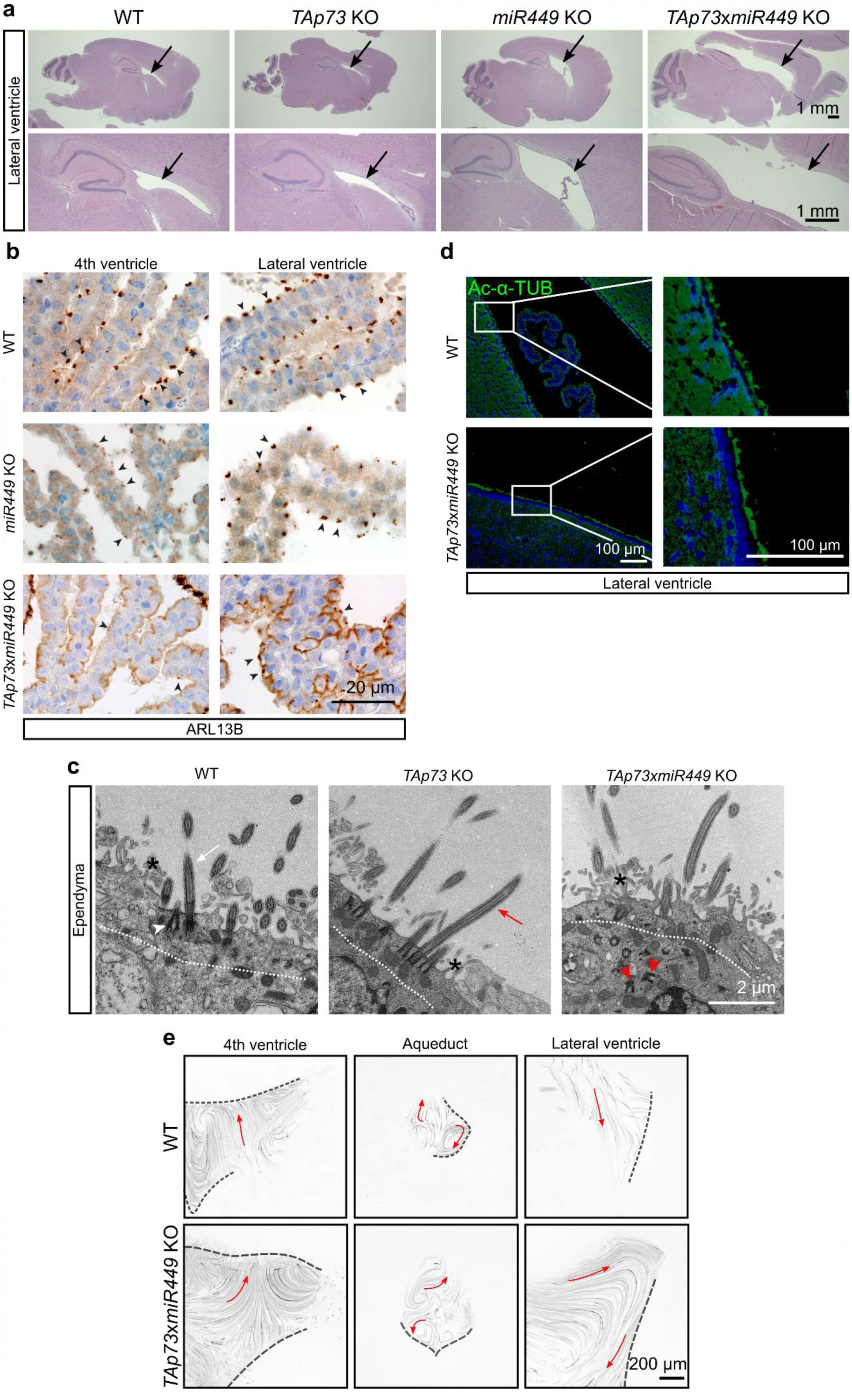
*miR449* collaborates with *TAp73* in brain multiciliogenesis. (**a**) Representative images of H&E staining of brain sections from WT, *TAp73* KO, *miR449* KO, and *TAp73xmiR449* KO animals. Notice that *TAp73xmiR449* KO mice display enlarged lateral ventricles (arrows). (**b**) Representative images of the expression of ARL13B in cilia (arrowheads) of CP in the 4^th^ and lateral ventricle from WT, *TAp73* KO, *miR449* KO, and *TAp73xmiR449* KO animals. (**c**) Representative photomicrographs of transmission electron microscopy (TEM) of ependymal cells from WT, *TAp73* KO, and *TAp73xmiR449* KO mice. Dotted lines mark apical region of the cells. Notice that the WT and *TAp73* KO cell possesses abundant cilia (arrows) and basal bodies (arrowhead) docked to the apical surface, whereas the *TAp73*x*miR449* KO cell has disorganized basal bodies (red arrowheads) located further away from the apical surface. Interspersed microvilli are marked with asterisks. (**d**) Expression of Ac-α-TUB (green) in ependymal cells of the lateral ventricle from WT and *TAp73xmiR449* KO mice. Boxed regions are magnified on the right. DAPI staining (blue) labels nuclei. (**e**) Analysis of the movement of fluorescent beads along the ventricular system from WT and *TAp73*x*miR449* KO mice. Images of maximum intensity projections of representative movies of the 4^th^ and lateral ventricle, and aqueduct are shown (*n*=2). Red arrows mark the direction of bead flow. Bracket lines delineate ependymal layer lining the ventricles. Refer to **Supplementary Video 3c** for examples of recording of ciliary beating.

**Supplementary Fig. 10.**
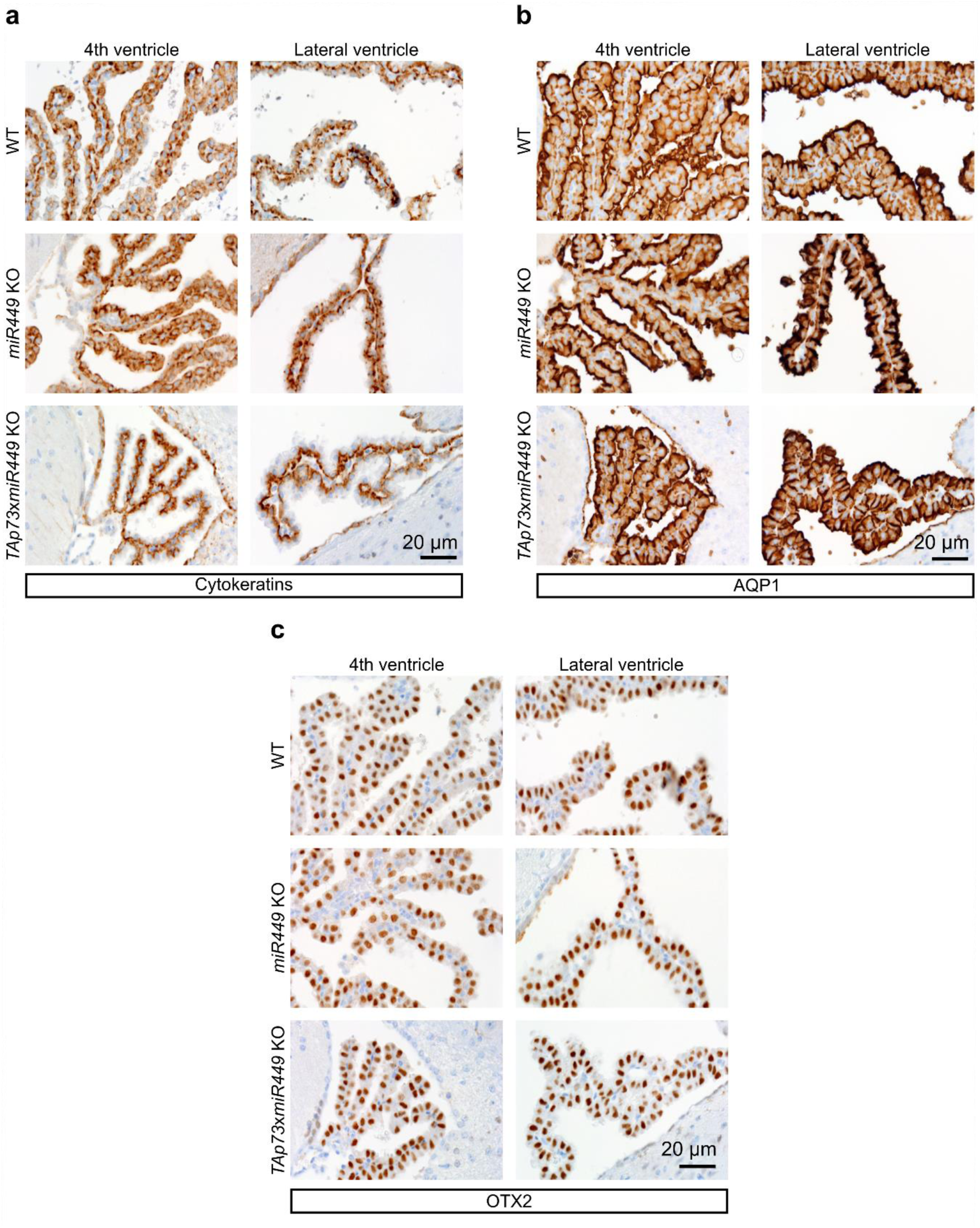
Combined loss of *TAp73* and *miR449* does not affect epithelial differentiation of choroid plexus cells. Representative images of the expression of cytokeratins (**a**), aquaporin 1 (AQP1, **b**), and orthodenticle homeobox 2 (OTX2, **c**) in CP epithelium of the 4^th^ and lateral ventricles from WT, *miR449* KO, and *TAp73xmiR449* KO mice.

**Supplementary Fig. 11.**
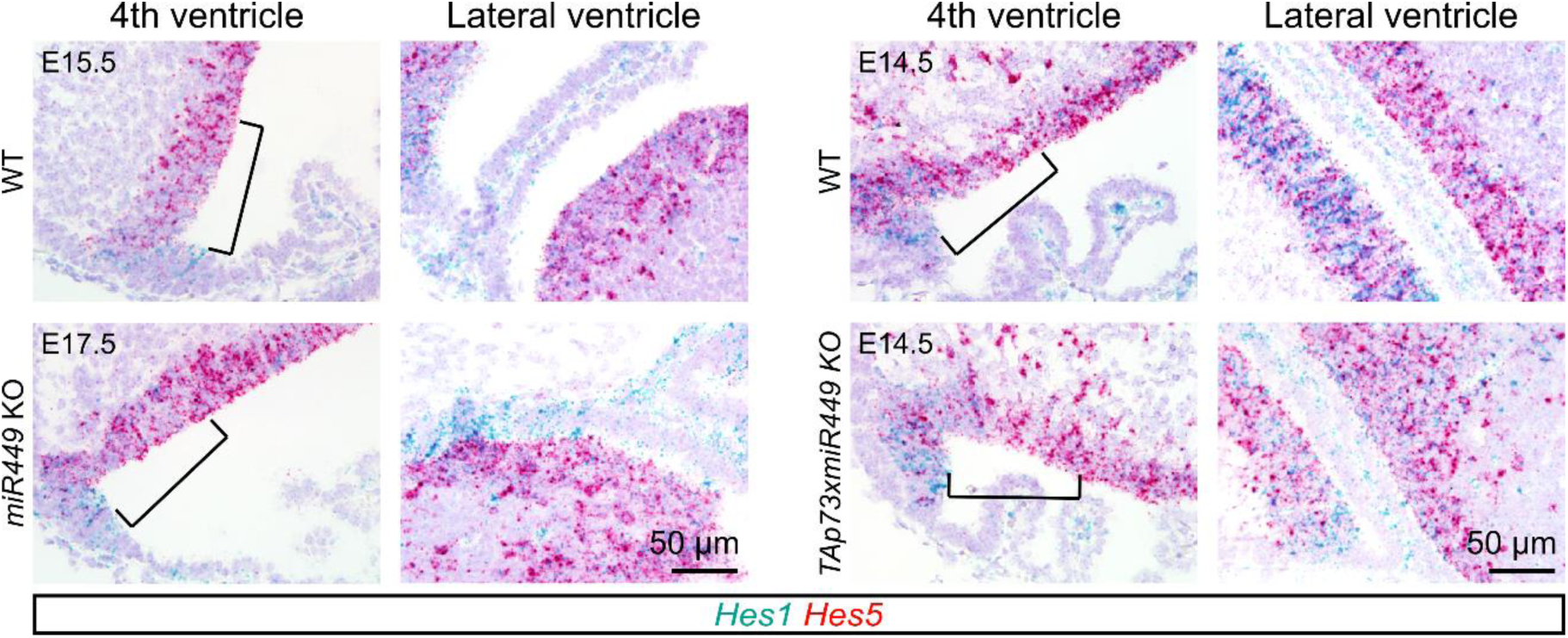
NOTCH signaling is unaltered in *miR449* KO and *TAp73xmiR449* KO brains. RNAscope analysis of the expression of NOTCH targets *Hes1* (blue) and *Hes5* (red) in roof plate of hindbrain and lateral ventricles from WT, *miR449* KO, and *TAp73xmiR449* KO mice.

**Supplementary Fig. 12.**
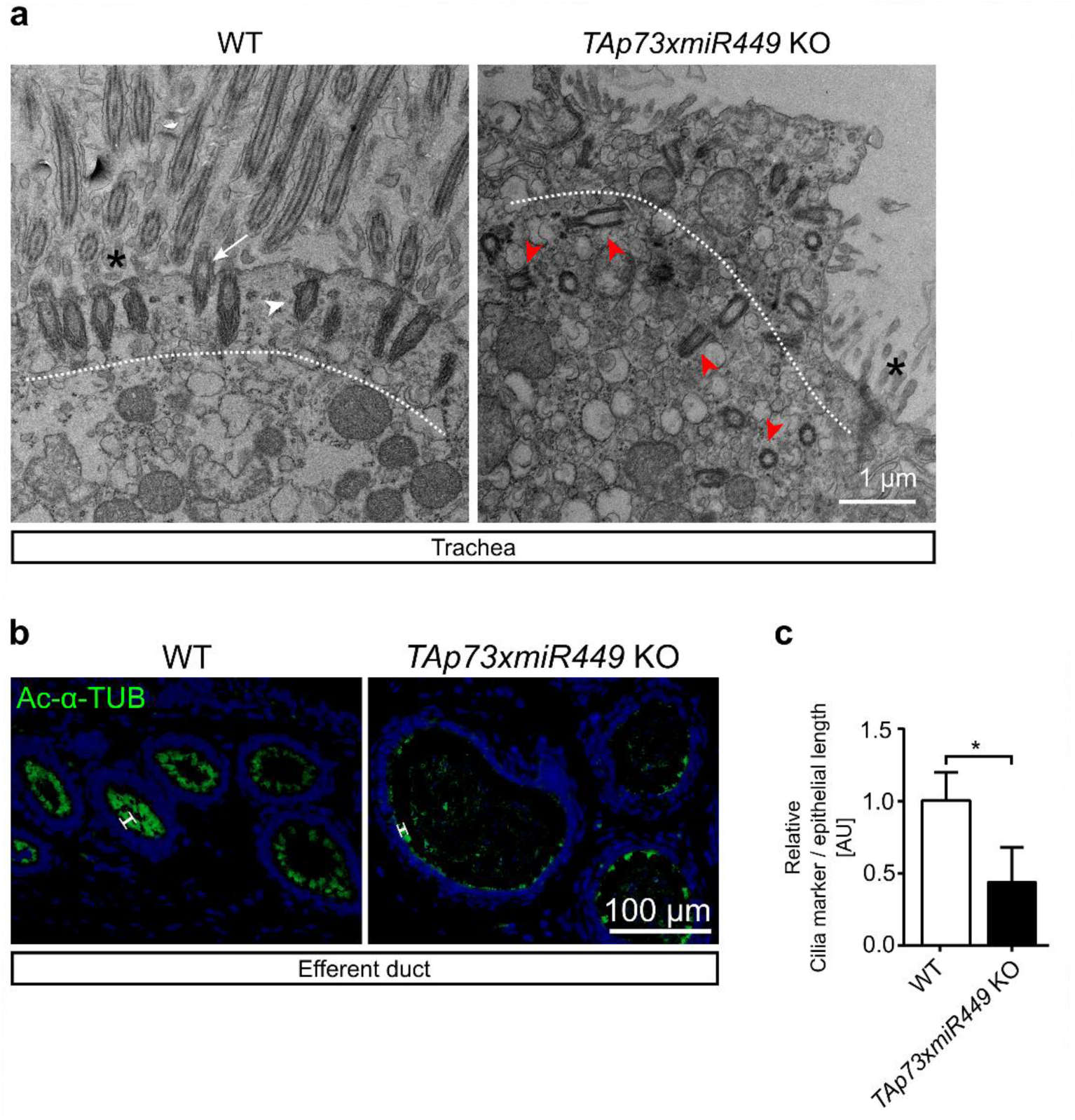
Additional loss of *miR449* does not exacerbate ciliary defects in the airways and efferent ducts in *TAp73* KO animals. (**a**) Representative photomicrographs of transmission electron microscopy (TEM) of trachea from WT and *TAp73xmiR449* KO mice. Dotted lines mark apical region of the cells. Notice the abundant cilia (white arrow) and clustered basal bodies (white arrowhead) docked to apical surface in the WT cell, whereas the mutant cell exhibits disorganized basal bodies (red arrowheads) located away from apical surface. Interspersed microvilli are marked with asterisks. (**b**) Representative images of the expression of Ac-α-TUB (green) in ED from WT and *TAp73xmiR449* KO mice. DAPI staining (blue) labels nuclei. Notice that mutant cells have less and shorter cilia (white bars) compared to WT mice. (**c**) Quantitation of Ac-α-TUB signals normalized to epithelial length (*n*=7 images from 4 WT mice; *n*=6 images from 3 *TAp73xmiR449* KO mice). Data are presented as mean ± SEM and relative to the WT group with \**P*<0.05.

### Supplementary Tables

**Supplementary Table 1.**
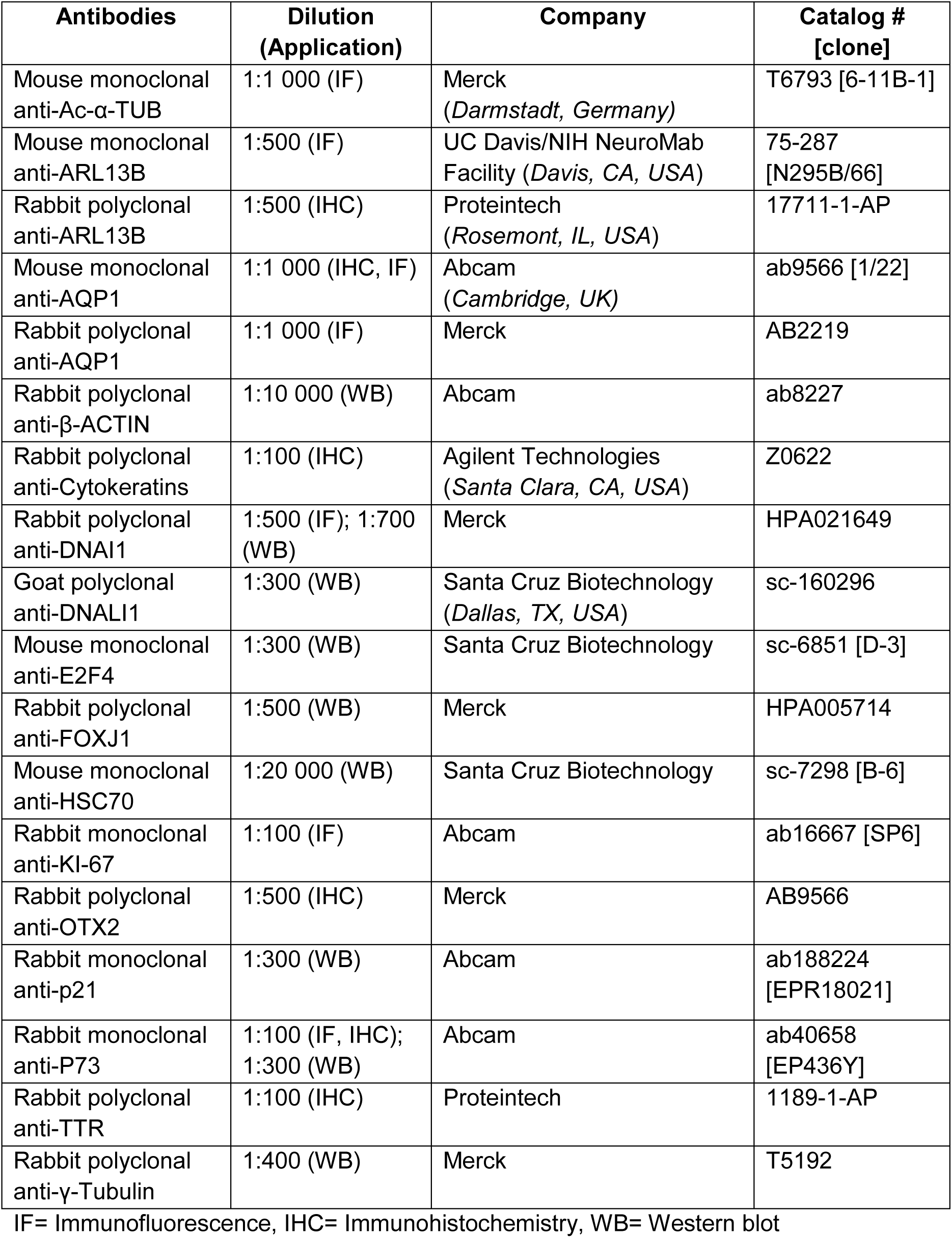
Primary antibody information.

**Supplementary Table 2.** Secondary antibody information.

| **Antibodies** | **Dilution**<br>**(Application)** | **Company** | **Catalog #** |
| --- | --- | --- | --- |
| Alexa Fluor 488 donkey anti-mouse | 1:500 (IF) | Invitrogen, Thermo Fisher Scientific (*Waltham, MA, USA*) | A21202 |
| Alexa Fluor 594 goat anti-rabbit | 1:500 (IF) | Invitrogen, Thermo Fisher Scientific | A11012 |
| Peroxidase-conjugated donkey anti-mouse | 1:10 000 (WB) | Jackson ImmunoResearch (*West Grove, PA, USA*) | 715-036-150 |
| Peroxidase-conjugated donkey anti-goat | 1:10 000 (WB) | Jackson ImmunoResearch | 705-036-147 |
| Peroxidase-conjugated donkey anti-rabbit | 1:10 000 (WB) | Jackson ImmunoResearch | 711-036-152 |
| Biotin-SP-conjugated AffiniPure Goat anti-Rabbit IgG | 1:1 000 (IHC) | Jackson ImmunoResearch | 111-065-144 |
| Biotin-SP-conjugated AffiniPure Donkey anti-Mouse | 1:1 000 (IHC) | Jackson ImmunoResearch | 111-065-144 |

IF= Immunofluorescence, IHC= Immunohistochemistry, WB= Western blot

**Supplementary Table 3.**
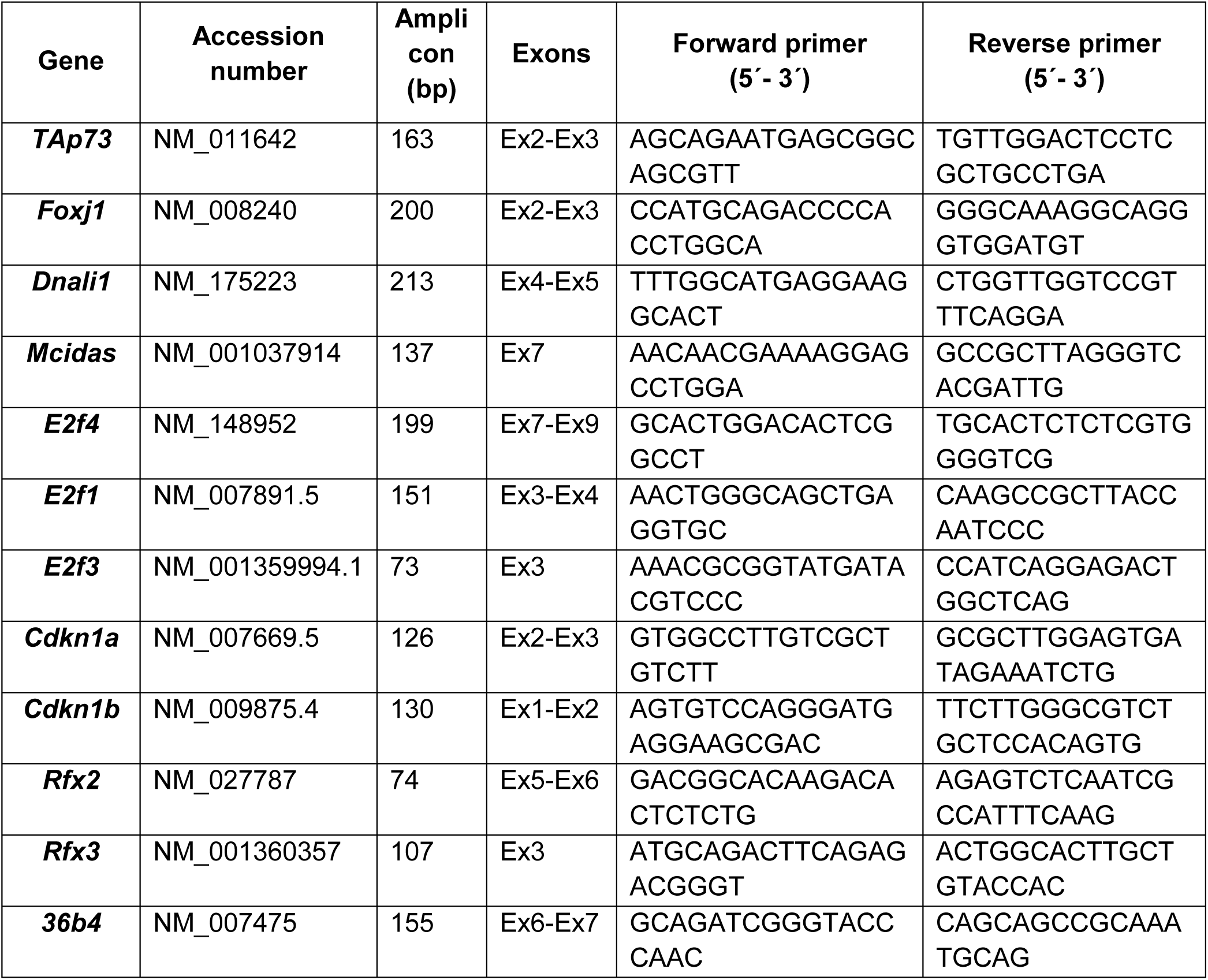
Sequence information for primers used in semi-quantitative PCR.

**Supplementary Table 4.**
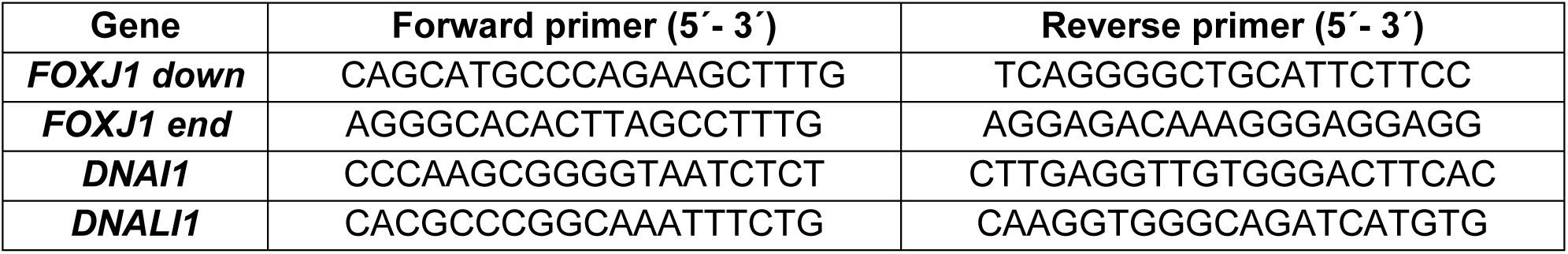
Sequence information for primers used in ChIP-qPCR.

**Supplementary Table 5.**
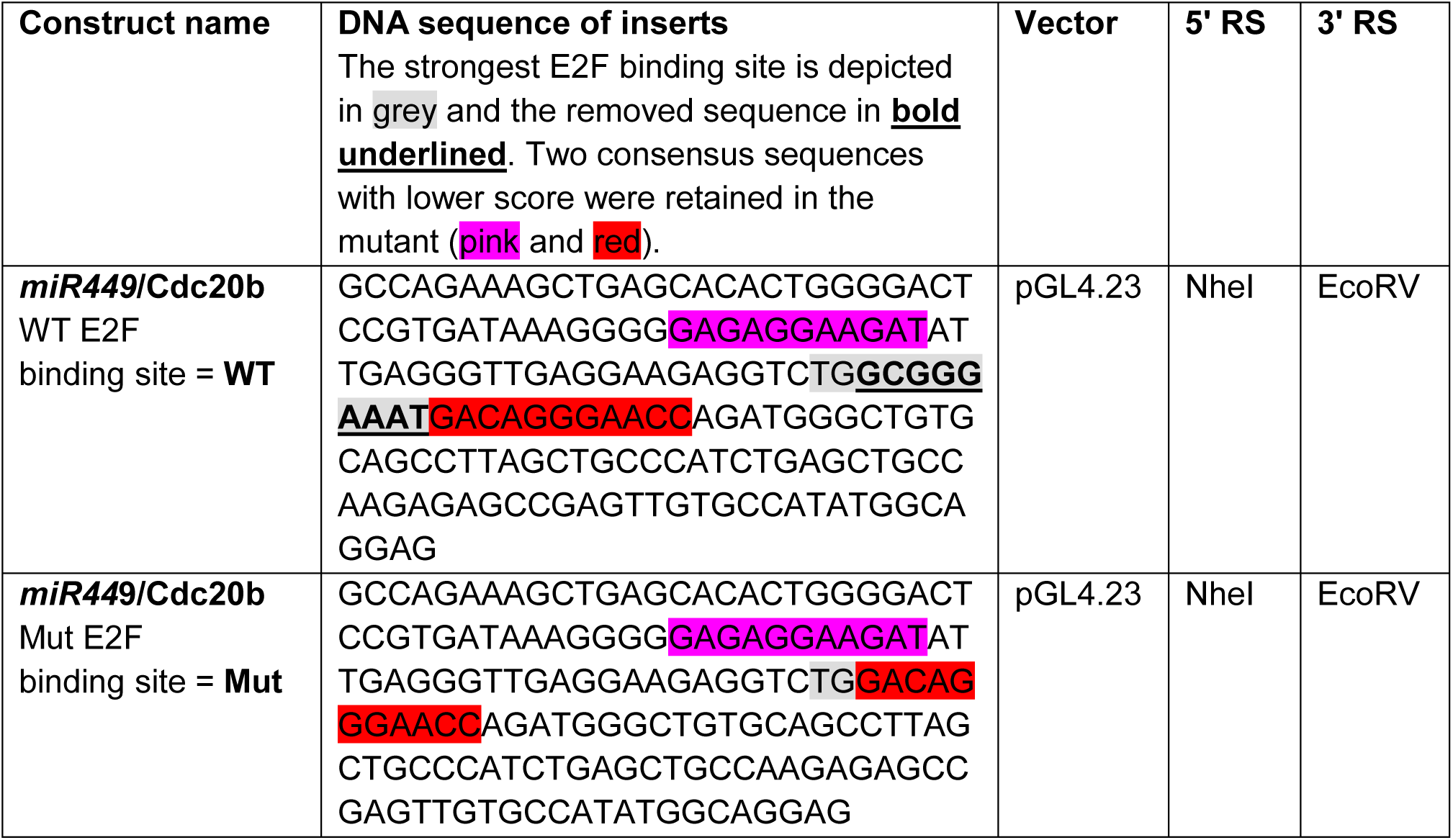
Luciferase constructs for E2F4/MCIDAS luciferase reporter assay. RS: Restriction site

**Supplementary Table 6.** Summary of small RNA-seq data from WT (*n*=3) and *TAp73* KO (*n*=4) brains. GEO accession number: **GSE108385.**

•small RNA-seq read counts from WT and TAp73 KO ventricles

•small RNA-seq differential gene expression results from WT vs. TAp73 KO ventricles.

### Supplementary Video legends

**Supplementary Video 1.** Movement of spermatozoa from *TAp73*^*+/-*^ (**a**, **b**) and *TAp73* KO mice (**c**, **d**).

**Supplementary Video 2.** Smooth muscle contraction in fallopian tubes from WT (**a**) and *TAp73* KO (**b**) mice.

**Supplementary Video 3.** Ciliary beating in WT (**a**, 3^rd^ ventricle), *TAp73* KO (**b**, lateral ventricle), and *TAp73xmiR449* KO (**c**, lateral ventricle) mice.

## References

1. Spassky N, Meunier A. The development and functions of multiciliated epithelia. Nat. Rev. Mol. Cell Biol. 2017;18:423–36.

2. Choksi SP, Lauter G, Swoboda P, Roy S. Switching on cilia: transcriptional networks regulating ciliogenesis. Dev. Camb. Engl. 2014;141:1427–41.

3. Terré B, Piergiovanni G, Segura-Bayona S, Gil-Gómez G, Youssef SA, Attolini CS-O, et al. GEMC1 is a critical regulator of multiciliated cell differentiation. EMBO J. 2016;35:942–60.

4. Arbi M, Pefani D-E, Kyrousi C, Lalioti M-E, Kalogeropoulou A, Papanastasiou AD, et al. GemC1 controls multiciliogenesis in the airway epithelium. EMBO Rep. 2016;17:400–13.

5. Zhou F, Narasimhan V, Shboul M, Chong YL, Reversade B, Roy S. Gmnc Is a Master Regulator of the Multiciliated Cell Differentiation Program. Curr. Biol. 2015;25:3267–73.

6. Boon M, Wallmeier J, Ma L, Loges NT, Jaspers M, Olbrich H, et al. MCIDAS mutations result in a mucociliary clearance disorder with reduced generation of multiple motile cilia. Nat. Commun. 2014;5:4418.

7. Ma L, Quigley I, Omran H, Kintner C. Multicilin drives centriole biogenesis via E2f proteins. Genes Dev. 2014;28:1461–71.

8. Stubbs JL, Vladar EK, Axelrod JD, Kintner C. Multicilin promotes centriole assembly and ciliogenesis during multiciliate cell differentiation. Nat. Cell Biol. 2012;14:140–7.

9. Danielian PS, Hess RA, Lees JA. E2f4 and E2f5 are essential for the development of the male reproductive system. Cell Cycle Georget. Tex 2016;15:250–60.

10. Danielian PS, Bender Kim CF, Caron AM, Vasile E, Bronson RT, Lees JA. E2f4 is required for normal development of the airway epithelium. Dev. Biol. 2007;305:564–76.

11. Marcet B, Chevalier B, Luxardi G, Coraux C, Zaragosi L-E, Cibois M, et al. Control of vertebrate multiciliogenesis by miR-449 through direct repression of the Delta/Notch pathway. Nat. Cell Biol. 2011;13:693–9.

12. Nemajerova A, Kramer D, Siller SS, Herr C, Shomroni O, Pena T, et al. TAp73 is a central transcriptional regulator of airway multiciliogenesis. Genes Dev. 2016;30:1300–12.

13. Marshall CB, Mays DJ, Beeler JS, Rosenbluth JM, Boyd KL, Santos Guasch GL, et al. p73 Is Required for Multiciliogenesis and Regulates the Foxj1-Associated Gene Network. Cell Rep. 2016;14:2289–300.

14. Blatt EN, Yan XH, Wuerffel MK, Hamilos DL, Brody SL. Forkhead transcription factor HFH-4 expression is temporally related to ciliogenesis. Am. J. Respir. Cell Mol. Biol. 1999;21:168–76.

15. Brody SL, Yan XH, Wuerffel MK, Song SK, Shapiro SD. Ciliogenesis and left-right axis defects in forkhead factor HFH-4-null mice. Am. J. Respir. Cell Mol. Biol. 2000;23:45–51.

16. Chen J, Knowles HJ, Hebert JL, Hackett BP. Mutation of the mouse hepatocyte nuclear factor/forkhead homologue 4 gene results in an absence of cilia and random left-right asymmetry. J. Clin. Invest. 1998;102:1077–82.

17. Yu X, Ng CP, Habacher H, Roy S. Foxj1 transcription factors are master regulators of the motile ciliogenic program. Nat. Genet. 2008;40:1445–53.

18. Crow J, Amso NN, Lewin J, Shaw RW. Morphology and ultrastructure of fallopian tube epithelium at different stages of the menstrual cycle and menopause. Hum. Reprod. Oxf. Engl. 1994;9:2224–33.

19. Lyons RA, Saridogan E, Djahanbakhch O. The reproductive significance of human Fallopian tube cilia. Hum. Reprod. Update 2006;12:363–72.

20. Raidt J, Werner C, Menchen T, Dougherty GW, Olbrich H, Loges NT, et al. Ciliary function and motor protein composition of human fallopian tubes. Hum. Reprod. Oxf. Engl. 2015;30:2871–80.

21. Vanaken GJ, Bassinet L, Boon M, Mani R, Honoré I, Papon J-F, et al. Infertility in an adult cohort with primary ciliary dyskinesia: phenotype–gene association. Eur. Respir. J. 2017;50:1700314.

22. Ilio KY, Hess RA. Structure and function of the ductuli efferentes: a review. Microsc. Res. Tech. 1994;29:432–67.

23. Lambot M-AH, Mendive F, Laurent P, Van Schoore G, Noël J-C, Vanderhaeghen P, et al. Three-dimensional reconstruction of efferent ducts in wild-type and Lgr4 knock-out mice. Anat. Rec. Hoboken NJ 2007 2009;292:595–603.

24. Hess RA. The Efferent Ductules: Structure and Functions [Internet]. In: Robaire B, Hinton BT, editors. The Epididymis: From Molecules to Clinical Practice. Boston, MA: Springer US; 2002 [cited 2017 Jan 19]. page 49–80. Available from: http://link.springer.com/10.1007/978-1-4615-0679-9_4

25. Hess RA. Small tubules, surprising discoveries: from efferent ductules in the turkey to the discovery that estrogen receptor alpha is essential for fertility in the male. Anim. Reprod. 2015;12:7–23.

26. Lun MP, Johnson MB, Broadbelt KG, Watanabe M, Kang Y-j., Chau KF, et al. Spatially Heterogeneous Choroid Plexus Transcriptomes Encode Positional Identity and Contribute to Regional CSF Production. J. Neurosci. 2015;35:4903–16.

27. Silva-Vargas V, Maldonado-Soto A, Mizrak D, Codega P, Doetsch F. Age-Dependent Niche Signals from the Choroid Plexus Regulate Adult Neural Stem Cells. Cell Stem Cell 2016;19:643–52.

28. Spassky N, Merkle FT, Flames N, Tramontin AD, García-Verdugo JM, Alvarez-Buylla A. Adult ependymal cells are postmitotic and are derived from radial glial cells during embryogenesis. J. Neurosci. Off. J. Soc. Neurosci. 2005;25:10–8.

29. Lun MP, Monuki ES, Lehtinen MK. Development and functions of the choroid plexuscerebrospinal fluid system. Nat. Rev. Neurosci. 2015;16:445–57.

30. Del Bigio MR. Ependymal cells: biology and pathology. Acta Neuropathol. (Berl.) 2010;119:55–73.

31. Li L, Grausam KB, Wang J, Lun MP, Ohli J, Lidov HGW, et al. Sonic Hedgehog promotes proliferation of Notch-dependent monociliated choroid plexus tumour cells. Nat. Cell Biol. 2016;18:418–30.

32. Tomasini R, Tsuchihara K, Wilhelm M, Fujitani M, Rufini A, Cheung CC, et al. TAp73 knockout shows genomic instability with infertility and tumor suppressor functions. Genes Dev. 2008;22:2677–91.

33. Song R, Walentek P, Sponer N, Klimke A, Lee JS, Dixon G, et al. miR-34/449 miRNAs are required for motile ciliogenesis by repressing cp110. Nature 2014;510:115–20.

34. Schindelin J, Arganda-Carreras I, Frise E, Kaynig V, Longair M, Pietzsch T, et al. Fiji: an open-source platform for biological-image analysis. Nat. Methods 2012;9:676–82.

35. Capece V, Garcia Vizcaino JC, Vidal R, Rahman R-U, Pena Centeno T, Shomroni O, et al. Oasis: online analysis of small RNA deep sequencing data. Bioinformatics 2015;31:2205–7.

36. Martin M. Cutadapt removes adapter sequences from high-throughput sequencing reads. EMBnet.journal 2011;17:10.

37. Dobin A, Davis CA, Schlesinger F, Drenkow J, Zaleski C, Jha S, et al. STAR: ultrafast universal RNA-seq aligner. Bioinforma. Oxf. Engl. 2013;29:15–21.

38. Liao Y, Smyth GK, Shi W. featureCounts: an efficient general purpose program for assigning sequence reads to genomic features. Bioinforma. Oxf. Engl. 2014;30:923–30.

39. Friedländer MR, Mackowiak SD, Li N, Chen W, Rajewsky N. miRDeep2 accurately identifies known and hundreds of novel microRNA genes in seven animal clades. Nucleic Acids Res. 2012;40:37–52.

40. Love MI, Huber W, Anders S. Moderated estimation of fold change and dispersion for RNA-seq data with DESeq2. Genome Biol. 2014;15:550.

41. Faubel R, Westendorf C, Bodenschatz E, Eichele G. Cilia-based flow network in the brain ventricles. Science 2016;353:176–8.

42. Holembowski L, Kramer D, Riedel D, Sordella R, Nemajerova A, Dobbelstein M, et al. TAp73 is essential for germ cell adhesion and maturation in testis. J. Cell Biol. 2014;204:1173–90.

43. Inoue S, Tomasini R, Rufini A, Elia AJ, Agostini M, Amelio I, et al. TAp73 is required for spermatogenesis and the maintenance of male fertility. Proc. Natl. Acad. Sci. 2014;111:1843–8.

44. Dacheux J-L, Dacheux F. New insights into epididymal function in relation to sperm maturation. Reproduction 2013;147:R27–42.

45. Mendive F, Laurent P, Van Schoore G, Skarnes W, Pochet R, Vassart G. Defective postnatal development of the male reproductive tract in LGR4 knockout mice. Dev. Biol. 2006;290:421–34.

46. Santos Guasch GL, Beeler JS, Marshall CB, Shaver TM, Sheng Q, Johnson KN, et al. p73 Is Required for Ovarian Follicle Development and Regulates a Gene Network Involved in Cell-to-Cell Adhesion. iScience 2018;8:236–49.

47. Huang X, Ketova T, Fleming JT, Wang H, Dey SK, Litingtung Y, et al. Sonic hedgehog signaling regulates a novel epithelial progenitor domain of the hindbrain choroid plexus. Dev. Camb. Engl. 2009;136:2535–43.

48. Kyrousi C, Lalioti M-E, Skavatsou E, Lygerou Z, Taraviras S. Mcidas and GemC1/Lynkeas specify embryonic radial glial cells. Neurogenesis 2016;3:e1172747.

49. Mori M, Hazan R, Danielian PS, Mahoney JE, Li H, Lu J, et al. Cytoplasmic E2f4 forms organizing centres for initiation of centriole amplification during multiciliogenesis. Nat. Commun. 2017;8:15857.

50. Caspary T, Larkins CE, Anderson KV. The Graded Response to Sonic Hedgehog Depends on Cilia Architecture. Dev. Cell 2007;12:767–78.

51. Lizé M, Herr C, Klimke A, Bals R, Dobbelstein M. MicroRNA-449a levels increase by several orders of magnitude during mucociliary differentiation of airway epithelia. Cell Cycle Georget. Tex 2010;9:4579–83.

52. Marcet B, Chevalier B, Coraux C, Kodjabachian L, Barbry P. MicroRNA-based silencing of Delta/Notch signaling promotes multiple cilia formation. Cell Cycle 2011;10:2858–64.

53. Otto T, Candido SV, Pilarz MS, Sicinska E, Bronson RT, Bowden M, et al. Cell cycletargeting microRNAs promote differentiation by enforcing cell-cycle exit. Proc. Natl. Acad. Sci. U. S. A. 2017;114:10660–5.

54. Redshaw N, Wheeler G, Hajihosseini MK, Dalmay T. microRNA-449 is a putative regulator of choroid plexus development and function. Brain Res. 2009;1250:20–6.

55. Yang X, Feng M, Jiang X, Wu Z, Li Z, Aau M, et al. miR-449a and miR-449b are direct transcriptional targets of E2F1 and negatively regulate pRb-E2F1 activity through a feedback loop by targeting CDK6 and CDC25A. Genes Dev. 2009;23:2388–93.

56. Lizé M, Pilarski S, Dobbelstein M. E2F1-inducible microRNA 449a/b suppresses cell proliferation and promotes apoptosis. Cell Death Differ. 2010;17:452–8.

57. Kyrousi C, Arbi M, Pilz G-A, Pefani D-E, Lalioti M-E, Ninkovic J, et al. Mcidas and GemC1 are key regulators for the generation of multiciliated ependymal cells in the adult neurogenic niche. Development 2015;142:3661–74.

58. Kim S, Ma L, Shokhirev MN, Quigley I, Kintner C. Multicilin and activated E2f4 induce multiciliated cell differentiation in primary fibroblasts. Sci. Rep. 2018;8:12369.

59. Ibanez-Tallon I. Dysfunction of axonemal dynein heavy chain Mdnah5 inhibits ependymal flow and reveals a novel mechanism for hydrocephalus formation. Hum. Mol. Genet. 2004;13:2133–41.

60. Banizs B, Pike MM, Millican CL, Ferguson WB, Komlosi P, Sheetz J, et al. Dysfunctional cilia lead to altered ependyma and choroid plexus function, and result in the formation of hydrocephalus. Dev. Camb. Engl. 2005;132:5329–39.

61. Banizs B, Komlosi P, Bevensee MO, Schwiebert EM, Bell PD, Yoder BK. Altered pH(i) regulation and Na(+)/HCO3(-) transporter activity in choroid plexus of cilia-defective Tg737(orpk) mutant mouse. Am. J. Physiol. Cell Physiol. 2007;292:C1409–1416.

62. Bill BR, Balciunas D, McCarra JA, Young ED, Xiong T, Spahn AM, et al. Development and Notch Signaling Requirements of the Zebrafish Choroid Plexus. PLoS ONE 2008;3:e3114.

63. Amirav I, Wallmeier J, Loges NT, Menchen T, Pennekamp P, Mussaffi H, et al. Systematic Analysis of CCNO Variants in a Defined Population: Implications for Clinical Phenotype and Differential Diagnosis. Hum. Mutat. 2016;37:396–405.

64. Guglielmino MR, Santonocito M, Vento M, Ragusa M, Barbagallo D, Borzì P, et al. TAp73 is downregulated in oocytes from women of advanced reproductive age. Cell Cycle Georget. Tex 2011;10:3253–6.

65. Hu W, Zheng T, Wang J. Regulation of Fertility by the p53 Family Members. Genes Cancer 2011;2:420–30.

66. Feng Z, Zhang C, Kang H-J, Sun Y, Wang H, Naqvi A, et al. Regulation of female reproduction by p53 and its family members. FASEB J. Off. Publ. Fed. Am. Soc. Exp. Biol. 2011;25:2245–55.

67. Fujitani M, Sato R, Yamashita T. Loss of p73 in ependymal cells during the perinatal period leads to aqueductal stenosis. Sci. Rep. 2017;7:12007.

68. Gonzalez-Cano L, Fuertes-Alvarez S, Robledinos-Anton N, Bizy A, Villena-Cortes A, Fariñas I, et al. p73 is required for ependymal cell maturation and neurogenic SVZ cytoarchitecture. Dev. Neurobiol. 2016;76:730–47.

69. Pefani D-E, Dimaki M, Spella M, Karantzelis N, Mitsiki E, Kyrousi C, et al. Idas, a Novel Phylogenetically Conserved Geminin-related Protein, Binds to Geminin and Is Required for Cell Cycle Progression. J. Biol. Chem. 2011;286:23234–46.

70. Bao J, Li D, Wang L, Wu J, Hu Y, Wang Z, et al. MicroRNA-449 and MicroRNA-34b/c Function Redundantly in Murine Testes by Targeting E2F Transcription Factor-Retinoblastoma Protein (E2F-pRb) Pathway. J. Biol. Chem. 2012;287:21686–98.

71. Fededa JP, Esk C, Mierzwa B, Stanyte R, Yuan S, Zheng H, et al. MicroRNA-34/449 controls mitotic spindle orientation during mammalian cortex development. EMBO J. 2016;35:2386–98.

72. Agostini M, Tucci P, Killick R, Candi E, Sayan BS, Rivetti di Val Cervo P, et al. Neuronal differentiation by TAp73 is mediated by microRNA-34a regulation of synaptic protein targets. Proc. Natl. Acad. Sci. U. S. A. 2011;108:21093–8.

73. Medina-Bolívar C, González-Arnay E, Talos F, González-Gómez M, Moll UM, Meyer G. Cortical hypoplasia and ventriculomegaly of p73-deficient mice: Developmental and adult analysis: p73 in developing and adult cortex. J. Comp. Neurol. 2014;522:2663–79.

74. Yang A, Walker N, Bronson R, Kaghad M, Oosterwegel M, Bonnin J, et al. p73-deficient mice have neurological, pheromonal and inflammatory defects but lack spontaneous tumours. Nature 2000;404:99–103.

75. Tissir F, Ravni A, Achouri Y, Riethmacher D, Meyer G, Goffinet AM. DeltaNp73 regulates neuronal survival in vivo. Proc. Natl. Acad. Sci. U. S. A. 2009;106:16871–6.

76. Koeppel M, van Heeringen SJ, Kramer D, Smeenk L, Janssen-Megens E, Hartmann M, et al. Crosstalk between c-Jun and TAp73alpha/beta contributes to the apoptosis-survival balance. Nucleic Acids Res. 2011;39:6069–85.

77. Diez-Roux G, Banfi S, Sultan M, Geffers L, Anand S, Rozado D, et al. A High-Resolution Anatomical Atlas of the Transcriptome in the Mouse Embryo. PLoS Biol. 2011;9:e1000582.

## References

1. Koeppel M, van Heeringen SJ, Kramer D, Smeenk L, Janssen-Megens E, Hartmann M, et al. Crosstalk between c-Jun and TAp73alpha/beta contributes to the apoptosis-survival balance. Nucleic Acids Res. 2011;39:6069–85.

